# Selectivity Dilution in Three-Factor Learning: A Finite-Population Model with Potential Implications for Motor Development and Post-Stroke Motor Rehabilitation

**DOI:** 10.64898/2026.10.14.759072

**Authors:** Zheng Liu, Chunyu Wu, Zhenhao Liu, Baohua Hu, Jingsong Mu

## Abstract

Motor development and post-stroke motor recovery may share underlying learning mechanisms, but the early mechanical environments of the two processes differ strikingly. During early development the fetus moves within amniotic fluid, where buoyancy substantially reduces gravitational load and enables spontaneous low-resistance movement; in contrast, post-stroke motor reconstruction occurs without this fluid low-resistance phase. Whether and how the absence of a fluid low-resistance phase influences post-stroke functional recovery is the central testable question raised by the selectivity dilution framework. Three-factor learning rules use broadcast neuromodulatory signals to convert eligibility traces into lasting synaptic changes. Because the signal is broadcast, it may reinforce all eligible synapses that are active during reinforcement without obvious deviation, not only those that caused the successful outcome. This framework offers a testable mechanistic principle for early motor learning and neurorehabilitation, and may inform the design of biologically plausible, non-backpropagation learning algorithms in artificial systems.

**Highlights:**

- Broader recruitment dilutes success-conditioned selectivity in finite populations.
- Left Edge and Dilution Zone formalize the success-selectivity trade-off.
- Even with high neural plasticity, post-stroke F-R mismatch is more likely to reinforce neural connections targeting anatomically torque-advantaged muscle groups.
- Dynamic resistance titration may restore F-R matching and selective reinforcement.

## Box 1.

**Key definitions**

### Eligible synapse number *n*

The number of synapses with active eligibility traces at the time of a phasic modulatory signal. In the computational model, *n* is the candidate pool for reinforcement.

### Global modulatory activation (GMA)

A dimensionless proxy for diffuse, non-selective neural drive, scaled by the number of co-activated neurons per time window. GMA is subserved by brainstem ascending reticular pathways (which also include the classic unspecific system neurons)^1^ and cortical top-down projections^2^ that enhance global excitability during the processing of incoming stimuli or effortful motor attempts ^3–6^. GMA is not identical to *n*; it influences *n* through multiple factors, perhaps including co-activation breadth, temporal window, oscillatory state, and inhibition, among others.

### Selectivity dilution

The reduction in the proportion of task-relevant synapses among reinforced synapses as *n* increases. It is quantified by the decay of *S*_*R*_(*n*).

### Sustainable reinforcement window (SRW)

The range of *n* where *P*_*success*_(*n*) > *θ*. Here *θ* = 0.1 is used for illustration, which means that on average, one out of every ten attempts succeeds. *θ* values significantly below this threshold make it unfavorable for animals to complete learning within an acceptable actual time frame, or to cope with spontaneous synaptic weakening.

### Left Edge / selective low-*n* regime

The subregion of the SRW where *S*_*R*_(*n*) > *ϕ*, with *ϕ* a task-specific threshold above the population baseline. Here reinforcement still confers a specificity advantage over random co-activation. Define the selective region *L*_*R*_ = {*n* : *P*_*R*_(*n*) > *θ* and *S*_*R*_(*n*) > *ϕ*_*sel*_}, with *ϕ*_*sel*_ = 0.05 used in the present analyses. The term “Left Edge” refers to the low-*n* selective interval only where such an interval has been verified.

### Dilution Zone / non-selective high-n regime

The subregion of the SRW where *S*_*R*_(*n*) ≤ *ϕ*. Reinforcement occurs, but with insufficient selectivity to complete the learning within an acceptably realistic timeframe. The dilution region is {*n* : *P*_*R*_(*n*) > *θ* and *S*_*R*_(*n*) ≤ *ϕ*_*sel*_}. It denotes lower selectivity under the chosen criterion.

### F-R matching

Alignment between residual voluntary force *F* and resistance *R*. Mismatch forces GMA upregulation to achieve movement. F-R matching is a proposed relation between measured voluntary torque capacity *C* and external torque threshold *R*. Distinguish *C* from the random trial torque *F*_*n*_. Mismatch may alter recruitment or cause failure.

## Introduction

Motor development and post-stroke motor recovery may share underlying learning mechanisms^7–13^, but the early mechanical environments of the two processes differ strikingly. During early development, the fetus moves within amniotic fluid, where buoyancy substantially reduces gravitational load and enables spontaneous low-resistance movement. Post-stroke motor reconstruction, by contrast, occurs without this fluid low-resistance phase. Whether and how the absence of a fluid low-resistance phase influences post-stroke functional recovery remains insufficiently examined, and constitutes one of the core motivations of the present work. Three-factor learning rules have been proposed as a framework for plasticity gated by neuromodulation. In contrast to two-factor Hebbian rules, three-factor rules introduce a third signal—such as dopamine or other neuromodulators—that converts transient synaptic eligibility traces into lasting weight changes^14–17^. For trial-based learning, as

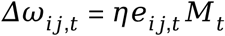

with the update interval specified, where *e*_*ij*_ is a transient eligibility trace and *Mt* is a broadcast modulatory signal. This scheme elegantly solves temporal credit assignment: synapses active shortly before reinforcement are consolidated^18–22^. Eligibility traces have been observed across striatum, cortex, and hippocampus^21,23,24^, with dopamine acting within a critical time window of approximately 0.3–2s^22,23^. Eligibility traces can bridge delays between activity and modulatory feedback, but do not by themselves identify the causal contribution of each synapse. VTA-M1 dopaminergic projections are necessary for skill acquisition^25–27^, with phasic release occurring preferentially on successful rather than failed trials^28–30^. Although the precise functional interpretation of these signals remains debated, the dilution constraint does not depend on resolving this debate—it arises from the broadcast nature of the modulatory signal itself. Studies support a role for dopaminergic projections to M1 in motor-skill acquisition.

Recent evidence suggests that DA signals in M1 are temporally linked to both movement execution and reward consumption (two phasic components)^30^. A short-latency DA signal^31^, occurring near movement onset, may reinforce the motor command and its associated eligibility traces. A subsequent DA signal, aligned with reward delivery, may reinforce motor representations or sensory cues that remain active at reinforcement. Both components are broadcasts^32^, and both reinforce whatever eligibility traces remain active at the time of release— regardless of whether those synapses contributed causally to the successful outcome. Here we focus on selectivity dilution in the first phasic stage. Movement- and reward-linked dopamine components may interact with temporally structured eligibility. We hypothesize that an early component can contribute to updating recently active motor pathways. The model assumes a common positive modulatory signal and equal positive eligibility within the sampled ensemble; these assumptions require experimental validation.

We propose that selectivity dilution is not a mere epiphenomenon but a systems-level constraint that may shape motor learning in health and may become maladaptive after stroke. The nervous system cannot neglect this trade-off: the breadth of activation required to overcome biomechanical resistance versus the precision of synaptic reinforcement. This framework applies primarily to exploratory attempts and early learning. In well-practiced, predictable movements with minimal phasic dopamine, this dilution effect may be negligible.

### 1. Selectivity dilution in the specified finite-population model

Consider a nascent network in which upstream representational units connect randomly and weakly to downstream motor outputs. On each trial, a subset of synapses acquires eligibility traces. When a broadcast modulatory signal arrives, the synapses with active eligibility traces form the candidate pool of size *n*. These traces are consolidated according to the three-factor rule. In this model, all sampled elements have equal positive eligibility and receive equal positive increments on successful trials. Under this assumption, the target fraction *r*_*n*_/*n* also equals the target share of the update.

Let *r* be the number of task-relevant synapses within this eligible pool of size *n*. We define selectivity as the conditional proportion of task-relevant synapses among eligible synapses on successful trials:

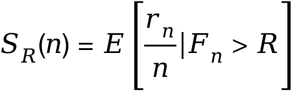

where *F*_*n*_ is net force and *R* is resistance. When *n* is small, success occurs only when the pool happens to be enriched in task-relevant synapses, so *S*_*R*_(*n*) is high. As *n* grows, the success condition becomes easier to satisfy, and *S*_*R*_(*n*) approaches the population baseline *p*_1_. This monotonic decline is the selectivity dilution constraint. Write *S*_*R*_(*n*) = *E*[*r*_*n*_/*n* | *F*_*n*_ > *R*] for *n* ≥ 1 and *P*_*R*_(*n*) > 0. For the specified extension model, *S*_*R*_(*n*) decreases over the verified domain even though *P*_*R*_(*n*) is non-monotone (Figure 1H,I). Success is eventually lost because the full-population torque is −600. In this extension model, as *n* increases, successful samples are increasingly constrained by the negative population mean and must draw from the upper tail of a distribution whose overall mean is negative. Conditional on success, the target fraction declines over the verified range because the success event becomes less informative about target enrichment as the sampling variance grows.

**Figure 1.**
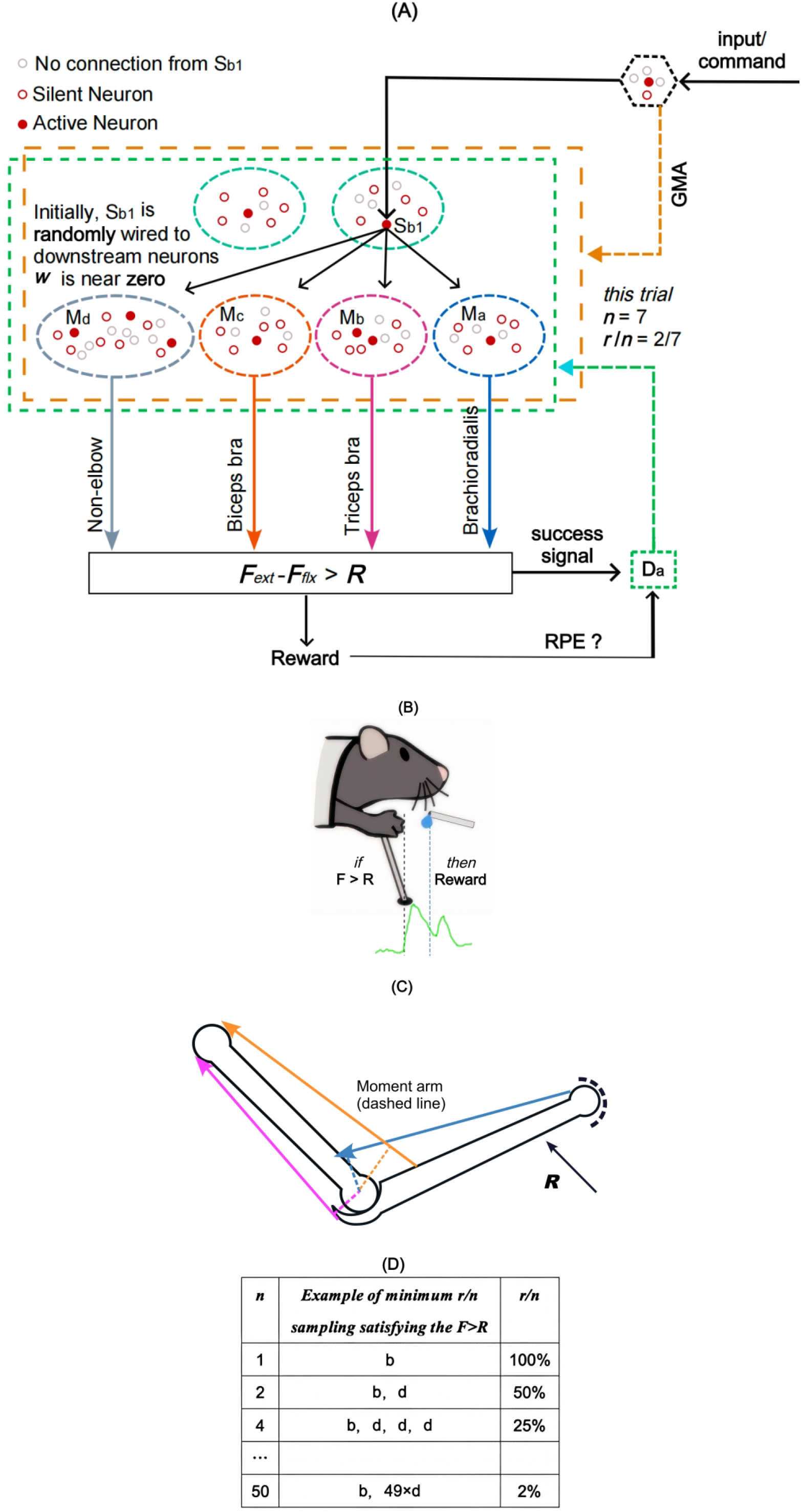
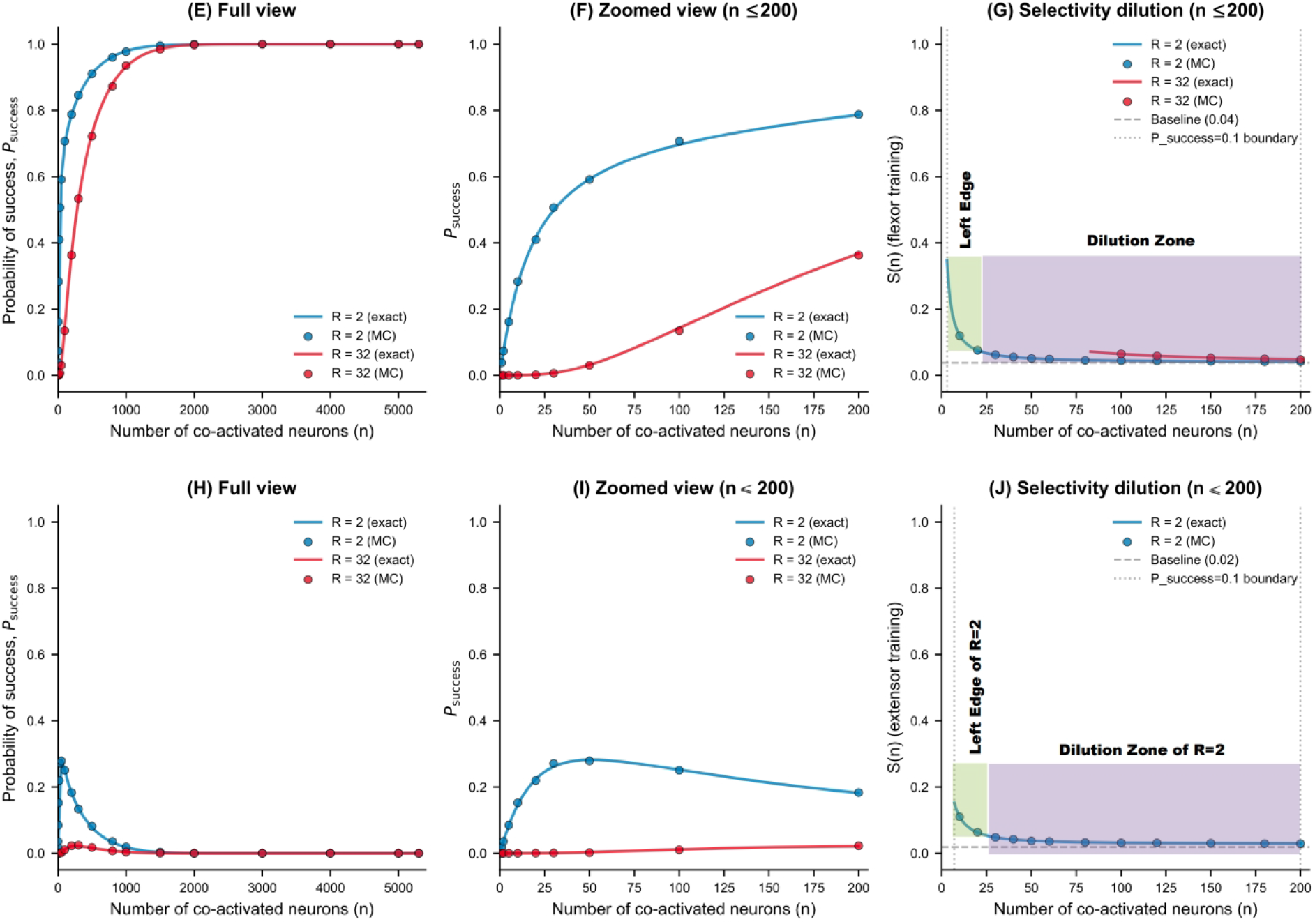
The selectivity dilution framework. (A–C) Conceptual diagrams; torque weights are illustrative and not estimated from the depicted moment arms. The dopamine trace in B is schematic. (D) Examples of minimum feasible target fractions for *R*=2, using one +9 element and otherwise zero-contribution elements. These examples are not estimates of *S*_*R*_(*n*). For example, at *n* = 50, *S*_*R*_(*n*) ≈ 0.03785, whereas the minimum illustrated fraction is 0.02. (E, F, H, I) Success probabilities for flexion (E, F) and extension (H, I). (G, J) Selectivity, displayed with an explicitly stated *ϕ*_*sel*_ = 0.05. Note that for R=32, SRW does not exist (J). Label n = 200 as the plot limit, not an SRW boundary. The data points presented here were obtained using the Monte Carlo method.

The system therefore faces a trade-off. A movement must recruit enough motor neurons (the number of eligible synapses increases accordingly) to generate the condition where *F*_*n*_ > *R*, but broader recruitment increases *n* and dilutes selectivity. We define the sustainable reinforcement window (SRW) as the range of *n* where

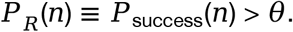

Within the SRW, the Left Edge is the subregion where *S*(*n*)> *ϕ*_sel_ reinforcement still confers a detectable specificity advantage over random co-activation, allowing learning to be completed in acceptable time. The Dilution Zone is the subregion where *S*_*R*_(*n*) ≤ *ϕ*_*sel*_, where reinforcement becomes lacking selectivity. A high resistance *R* narrows the SRW and can even completely eliminate the Left Edge (Figure 1G, J, red curve).

Two pathological regimes should be distinguished. First, the SRW persists but the entire region becomes a Dilution Zone, so the Left Edge disappears (Figure 1G, red curve. Note that this example uses elbow flexion as the case). In the second, the SRW itself is abolished because *P*_*success*_(*n*) < *θ* for all *n* (Figure 1H, I, red curve). Both are maladaptive, but they arise under different mechanical conditions. The first case strengthens the pathways of muscle groups with an anatomical advantage in output torque, which is reminiscent of the common flexor synergy after stroke and the lower limb extensor synergy. Figure 1 illustrates this logic qualitatively, and Figure 2 illustrates the dynamic changes of downstream circuit connections. In a flexor-dominant system, extensor-targeted movements exhibit a non-monotonic relationship between *n* and *P*_*success*_(*n*): low *n* fails to generate sufficient force, intermediate *n* allows variance-driven success, and high *n* is dominated by flexor bias^33,34^. Low resistance permits operation at the Left Edge, where *n* is small and *S*_*R*_(*n*) is sufficient. High resistance forces operation at high *n*, driving the system into the Dilution Zone or even lacking SRW. Exact derivations and numerical evaluations are provided in SI 3–4. In extension with *θ* = 0.1, the exact windows are 7–410 for *R* = 2 and 7– 299 for *R* = 8; no window exists for *R* = 32. These classifications are operational model results and do not by themselves establish maladaptive learning (see SI 3.5 and Figure SI3).

**Figure 2.**
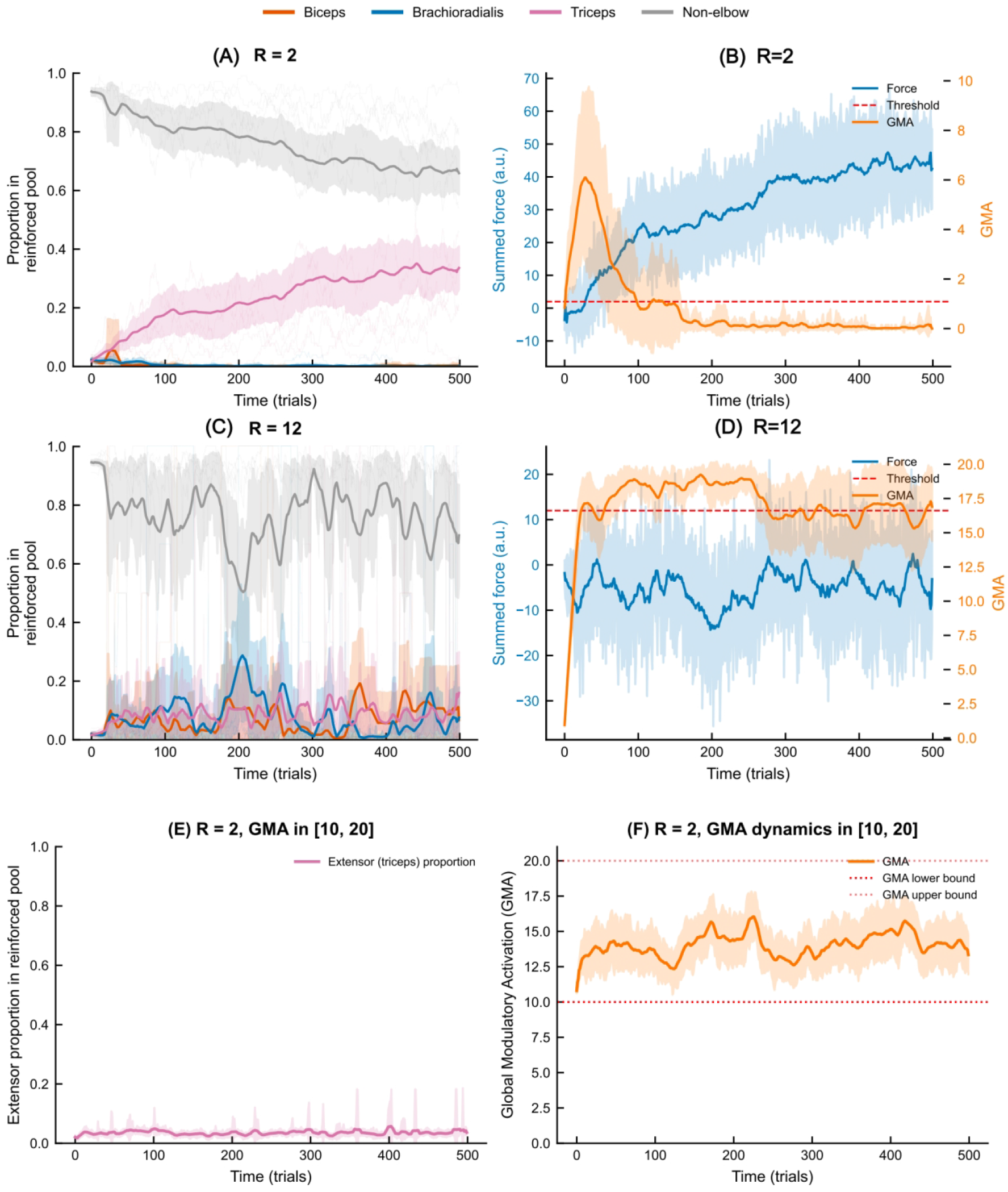
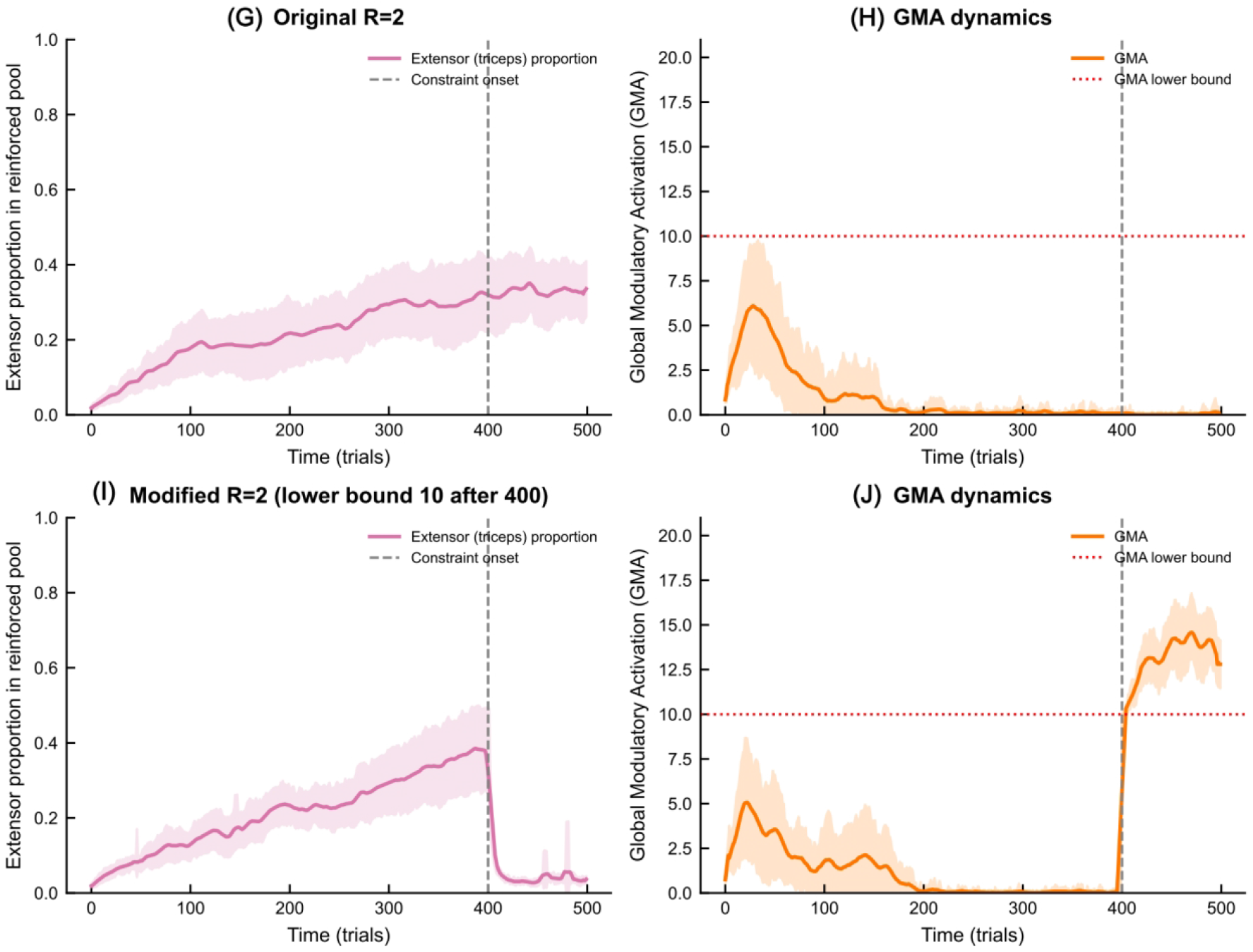
Illustrative simulations of the dynamic ensemble model. (A–B) Composition of the reinforced pool and force/GMA dynamics for R = 2. (C – D) For R = 12, the proportion of connections to task-relevant pathways did not increase, whereas the proportion of connections to non-task-relevant pathways did not significantly decrease. (E–F) Forced high GMA impairs an established extensor pathway. (G–H) The proportion of connections to task-relevant pathways gradually increases, while it degenerates when the modified condition enforces GMA ≥ 10 after trial 400 (I–J). The model is specified in SI Section 5. Twenty independent runs per condition, seed 32, 500 trials.

As mentioned in this model, randomly initialized network connections are adopted, and their connection strengths are set to zero. This framework may also circumvent a long-standing debate regarding whether sensory or motor drives initiate network refinement. Sensory stimulation is necessary, but it does not initially need to directly drive the network output. At this stage, in response to sensory stimuli, the nascent network can upregulate GMA through the ascending activation pathway after several attempts, which increases the variance of torque output to generate a success signal (Figure 1A). Specify how exploratory output is generated before learned connections become effective. If learned S_b_-to-output weights start at zero, a separate background drive must produce the fixed output contributions used in the sampling model.

### 2. Anatomical and physiological substrates

The dilution constraint is grounded in the anatomy of dopaminergic and diffuse modulatory systems. Midbrain dopamine neurons project diffusely to target structures. When dopamine is released—whether in response to unexpected reward, salient sensory events, or successful movement outcomes—it reaches a vast territory of synaptic terminals simultaneously^32^. VTA dopamine neurons may converge on a common reward prediction error signal^35^, while tail-of-striatum dopamine can encode a value-free action prediction error^36^. Both are broadcast teaching signals. Both reinforce whatever eligibility traces remain active at the time of release. Dopamine signals exhibit heterogeneity across neurons, projections, and behavioral variables, including specialized coding reported by Engelhard et al^35^. We approximate the signal within the modeled ensemble as shared, while recognizing that local release, receptors, and eligibility can confer additional specificity.

GMA is not a single anatomical pathway. It is subserved by at least two components. The first is the brainstem ascending reticular system, which sets a state-dependent excitability baseline^1^. The second consists of cortical top-down projections from dorsal premotor cortex^2^. These projections add phasic excitability during novel, complex, or effortful motor attempts, even without high mechanical load. Thus, excessive GMA can arise from mechanical necessity, cognitive effort, or both. Descending control of force is not exclusively corticospinal: both the corticospinal and reticulospinal tracts contribute to the force of contraction^37^, pontine reticular activity scales with handgrip force in humans^38^, and resistance-training tempo can selectively modulate corticospinal versus reticulospinal excitability^39^. Increases in premotor corticospinal excitability are themselves largely spinally mediated^40^. Ascending arousal systems and premotor-to-M1 interactions are candidate contributors to the drive summarized by GMA.

Several mechanisms may enhance specificity. Feedback-based gating can tag synapses responsible for a selected action^41–43^. Thalamocortical disinhibition can gate long-term potentiation^44^. Neural oscillations concentrate firing into Up states, which may reduce irrelevant eligibility traces during Down states. A narrower phasic dopamine window may constrain reinforcement to synapses activated within a precise interval^22,23^. These mechanisms may alleviate dilution, but they do not eliminate it, especially when F-R mismatch forces chronically high GMA. In the specified model, selectivity dilution is a recruitment-dependent constraint that arises when broadcast reinforcement acts on a uniformly sampled eligible ensemble, not a failure of any particular gating mechanism.

### 3. F-R mismatch and maladaptive reinforcement

In healthy individuals operating within biomechanical capacity, the system can maintain *n* near the Left Edge. Movement success is achieved with moderate GMA, allowing selective strengthening of task-relevant pathways. Diffuse projection systems (including classic unspecific system neurons) may serve a necessary function here: they provide the exploratory variability necessary for motor learning^45^. We hypothesize that efficient learned recruitment can achieve success with a smaller eligible ensemble.

After corticospinal tract damage, the residual voluntary force *F* may fall far below the environmental resistance *R*. The system then faces a stark choice: fail to move, or upregulate GMA to increase *n* until *P*_*success*_ > *θ*. If *n* is upregulated into the Dilution Zone, the system may achieve movement, but at the cost of reinforcing a broad, lacking-selective set of connections. Under sufficiently high resistance, even the SRW itself may be abolished (Figure 1J).

This F-R mismatch can abstractly create a maladaptive positive feedback loop. High GMA reduces reinforcement selectivity, causing non-target pathways—especially anatomically dominant pathways—to be repeatedly co-strengthened. Because flexor pathways are anatomically dominant in the upper limb^33,34^, non-selective reinforcement is more likely to consolidate these pathways over repeated trials and even generate negative force, which represents flexion force in this context (Figure 2D). This drives the system toward a low-recovery steady state that may clinically manifest as flexion synergy. Similarly, lower-limb training is more likely to strengthen the triceps surae muscle group, as this muscle group inherently has a greater anatomical advantage than the ankle dorsiflexion muscle group. Consistent with this, patients with moderate-to-severe impairment often quickly plateau: beyond a severity threshold, conventional rehabilitation cannot drive selective reorganization regardless of training dose^46^. This framework predicts that the system may become trapped not only because plasticity has ceased, but also because the reinforcement that occurs is maladaptive.

Existing clinical and neurophysiological observations are compatible with this framework. In a longitudinal fMRI study of hemiparetic stroke patients, Feydy and colleagues distinguished progressive focusing of activation toward contralateral sensorimotor cortex from persistent recruitment of widespread bilateral networks^47^. Persistent recruitment was associated with damage to M1 itself, whereas patients with spared M1 predominantly showed progressive focusing. This dissociation suggests that M1 integrity may determine whether the system can lower GMA and exit the Dilution Zone. When M1 is badly compromised, diffuse, non-selective recruitment may persist, locking the system into a state where selective reinforcement of alternative pathways remains difficult. Persistent widespread activation is compatible with several mechanisms, including altered recruitment after injury. It is not a direct measure of *n*, GMA, or *S*_*R*_(*n*). Relating these imaging findings to the dilution hypothesis requires within-subject measurements and control for lesion location and severity.

This framework does not deny the role of loss of central inhibition or other classical mechanisms of spasticity^48^. Rather, it offers a complementary computational mechanism: reticulospinal bias may be exacerbated by maladaptive reinforcement under gravitational loading, potentially contributing to progressive consolidation of flexor synergy in chronic stroke^49^. Consistent with this observation, shoulder abduction loading progressively reduces elbow extension range^50,51^ of motion and impairs volitional hand opening in a load-dependent manner^52^—these patterns align with the prediction that proximal sub-actions (e.g. shoulder abduction) requiring higher GMA can dilute distal neural selectivity.

Dilution is not inherently maladaptive. When existing low-GMA pathways cannot meet novel task demands, moderate GMA upregulation can broaden co-activation and transiently randomize network activity, serving an exploratory role. Early after stroke, transient increases in connectivity—such as contralateral-to-ipsilateral M1 coupling at 2 weeks—are consistent with a more randomised configuration^53^. With recovery, both decreased and increased connectivity tend to normalize toward healthy levels^53–55^. In severe cases, however, reduced connectivity between ipsilesional hemisphere and cerebellum, and between ipsilesional M1 and SMA, may persist^54^; consistent with this, acute patients with greater deficits show shorter high-centrality dwell time in ipsilesional ventral central sulcus and fewer dynamic interactions with contralateral central sulcus, SMA, and ipsilesional cerebellum^56^. Thus, early randomization may support exploration, whereas failure to normalize may reflect trapping in the Dilution Zone. Aphasia recovery similarly depends on dynamic reorganization^57^, suggesting that broader activation can expand the search space for compensatory pathways when core networks are compromised. These observations remain speculative. Dilution is therefore a double-edged sword: it may enable exploration, yet become maladaptive if the system remains trapped in the Dilution Zone. The therapeutic challenge is not to abolish dilution but to keep exploratory GMA increases transient, allowing a return to the Left Edge where selective reinforcement can consolidate task-relevant pathways.

### 4. Development and rehabilitation through the F-R lens

The framework offers a unified, testable interpretation of early sensorimotor development and post-stroke motor impairment^7–13^. During prenatal development, the fetus moves in amniotic fluid, an environment often described as reducing gravitational load and providing velocity-dependent resistance. Whether this approximates a form of dynamic F-R matching remains a hypothesis. It could permit nascent motor systems to achieve occasional success through variance-driven fluctuations in initially weak and diffuse output, with n maintained near the Left Edge. Over repeated cycles, effective connections would strengthen, the rudiment of the sensory-motor loop emerges (this provides a structural foundation for the subsequent rise of voluntary torque, as the PFC may innervate these types of loops), and GMA can progressively decrease while success probability is maintained.

This developmental interpretation is speculative. Fetal motor behavior is also shaped by endogenous spinal rhythms, sensory feedback, and neuromuscular maturation^58–60^. The framework does not claim that F-R matching is the sole teacher of early development. It proposes that mechanical matching provides a plausible additional constraint that may help explain the transition from diffuse, multi-joint fetal movements to focused, coordinated actions^58,61^. We believe that any species possessing fine motor skills in adulthood but that does not require excessive activity in low-resistance (including gravitational) environments during its early developmental stage can serve as a counterexample to test this hypothesis. A comparative test would require prespecified measures of developmental loading, activity, muscle capacity, and later motor precision, with controls for species-specific maturation and recruitment. Such evidence could test the hypothesis but is not a direct counterexample to the finite-population calculation. After stroke, we hypothesize that the same logic may apply in reverse. Corticospinal damage may reduce voluntary extensor torque far below gravitational resistance. To complete task-related movements, the system may upregulate GMA into the Dilution Zone; the resulting non-selective reinforcement would then be expected to consolidate circuits with anatomical advantages (such as the flexor muscle groups of the upper limbs and the triceps surae of the lower limbs). When stroke disrupts dominant low-threshold pathways of a mature network, sudden F-R mismatch may trigger pathological GMA upregulation. This process may reactivate previously disused higher-threshold pathways, and may clinically present as the re-emergence of primitive reflexes^62^ alongside diffuse, non-selective motor output. This pattern aligns with network-level randomization documented in longitudinal stroke connectivity studies^55,63,64^. It also aligns with the principle that recovery after stroke recapitulates ontogeny^7,8^. We propose that injury-related changes in recruitment may interact with learning under load. Reactivation of alternative pathways, primitive reflexes, and developmental parallels are possible interpretations requiring separate anatomical and longitudinal evidence; they are only inspired by the static dilution model or the present dynamic simulations.

This framework also explains why mild and moderate-to-severe patients may require different training environments. In mild impairment, residual force F remains close to gravitational resistance R, so a moderate increase in GMA can bring the system into the sustainable reinforcement window while keeping the eligible ensemble small enough to preserve selectivity; such patients can operate near the Left Edge even under full gravity. In moderate-to-severe impairment, F falls far below R, requiring a large GMA increase that expands n into the Dilution Zone or even abolishes the SRW. The former result is that broad, non-selective pathway connections with anatomical mechanical advantages are reinforced, while the latter result is no obvious recovery. Reducing external resistance restores F-R matching, returns the system toward the Left Edge, and allows selective reinforcement to resume. The model therefore predicts that low-resistance training is most necessary when residual force is far below the imposed load, and least necessary when the two are already matched.

The Left Edge strategy follows directly: dynamically match external resistance to the network’s current force output capacity. In rehabilitation, this means continuous, closed-loop titration of resistance rather than fixed high-load training. Long-term aquatic therapy, robotic exoskeletons, and sensorized devices with adaptive resistance schedules are candidate implementations. The principle is not to abolish effort, but to keep effort within a range where successful trials remain frequent enough to drive reinforcement while *n* remains small enough to preserve selectivity. Adaptive resistance is a candidate intervention suggested by the hypothesis. Its effects should be compared with dose- and success-matched control protocols in prospective studies.

Potential relevance to artificial learning systems. Beyond its implications for motor development and stroke rehabilitation, the present framework may also be relevant to artificial learning systems. Three-factor learning rules constitute a biologically plausible form of credit assignment that does not rely on backpropagation, and the selectivity dilution described here identifies a general trade-off between recruitment breadth and update specificity. The Left Edge strategy—dynamically constraining the eligible ensemble to maintain a high signal-to-noise ratio—can be viewed as a resource-constrained optimization principle that may inspire the update rules of connections in artificial neural networks and brain–computer interfaces.

### 5. Testable predictions

#### Prediction 1: TMS recruitment curve shifts

Among patients who successfully overcome flexion synergy, transcranial magnetic stimulation should reveal a leftward shift of the recruitment curve for elbow extensor muscles, while non-responders show no change or paradoxical flexor facilitation^65^. Test whether changes in a prespecified extensor TMS input-output measure correlate with independently measured recruitment selectivity and functional recovery.

#### Prediction 2: Pharmacological interaction

Dopaminergic agents such as levodopa may enhance motor recovery in patients who have already achieved F-R matching, but may be ineffective or even consolidate maladaptive circuits in patients who have not. This prediction appears to be consistent with the mixed results of dopamine-augmented rehabilitation trials: positive effects in mildly impaired patients who could complete motor assessments^66,67^, and null results in moderate-to-severe populations^68,69^. The recent negative ESTREL trial^70^ does not preclude benefit in the subset who had already achieved F-R matching at enrollment. Testing this prediction requires stratification by validated F-R matching criteria. For example, the ability to extend fingers when the shoulder is abducted may be explored as a candidate measure.

#### Prediction 3: Time-dependence of low-resistance exposure

For patients with moderate-to-severe impairment, low-resistance exposure should be most effective during early recovery. Increasing successful attempts per unit time or extending the effective recovery window may partially compensate for mechanical unloading by increasing the total number of stochastic successes within the critical period. The hypothesis predicts a possible interaction between loading, trial count, and recovery stage, but the present model contains no critical-period dynamics. More attempts can increase expected successes at fixed *P*_*R*_(*n*); they do not necessarily reproduce the selectivity or clinical effects of mechanical unloading.

#### Prediction 4: Residual co-activation as a signature of dilution

Even in well-optimized motor circuits, forceful isolated movements may evoke detectable co-activation of non-target motor pools. This phenomenon should be more pronounced in patients with impaired motor function where GMA is excessively upregulated. High-density EMG or functional MRI with motor tasks could test this prediction. Test whether non-target co-activation predicts subsequent task-specific learning after controlling for force, stability demands, and impairment. EMG and fMRI provide indirect recruitment measures.

#### Prediction 5: Progressive resistance calibration

Gradual, not step-like, increases in resistance should be essential for maintaining selectivity. Abrupt increases recruit both high- and low-threshold circuits together, diluting selectivity and ultimately increasing the randomization of network connections. This principle can be tested by comparing learning outcomes under gradually versus abruptly increasing task difficulty, or by using robotic exoskeletons programmed with adaptive resistance schedules. We hypothesize that gradual and abrupt resistance schedules may produce different recruitment and learning outcomes. Compare them under matched total practice, loading, and success exposure.

## Outstanding questions

1. Can the predicted non-monotonic relationship between operational proxies of GMA and *P*_*success*_ for extensor-targeted movements be tested in vivo during prespecified graded loading?
2. Does treatment response vary with prospectively measured recruitment and F-R matching, and does it exhibit an inverted-U relationship that holds true across alternative conditions? In other words, the response is optimal when matching is optimal, while performance declines under both overloading and underloading conditions.
3. Can low-resistance environments—aquatic or device-assisted—produce selective extensor strengthening in randomized controlled trials using prespecified EMG and kinematic outcome measures?
4. Does a validated, prespecified F-R matching measure explain recovery beyond baseline impairment and corticospinal integrity, after accounting for bounded scales and statistical coupling?

## Concluding remarks

The specified finite-population model exhibits a recruitment-dependent decline in success-conditioned target fraction over the verified parameter range. This motivates hypotheses connecting recruitment, resistance, and learning specificity. Generalization to structured neural circuits and clinical recovery requires explicit learning dynamics and experimental validation. Adaptive resistance is a testable intervention concept.

## Acknowledgments

The authors thank the rehabilitation team at The First Affiliated Hospital of the University of Science and Technology of China for their support in discussions. We also express our gratitude to Professor Chaomin Ni, Professor Ming Wu, and Professor Yan Huang (Department of Rehabilitation Medicine, the First Affiliated Hospital of USTC), Professor Quan Zhao, Professor Shucheng Yu (Department of Applied Mathematics, University of Science and Technology of China), and Ping Zhao (School of Mechanical Engineering, Hefei University of Technology) for their valuable assistance and discussions.

## Code and data availability

All simulation code, random seeds, parameter files, and plotting data for Figure 2 and SI Figure 1 are available in the Science Data Bank (ScienceDB) at https://doi.org/10.57760/sciencedb.015aw. The static exact combinatorial calculations are provided as reproducible scripts with arbitrary-precision integer comparisons.

## Supplementary Information

for “Selectivity Dilution in Three-Factor Learning: A Finite-Population Model with Potential Implications for Motor Development and Post-Stroke Motor Rehabilitation”

### SI 1. Where does the diffuse drive that elevates GMA originate?

The concept of GMA is anchored in a dual neuroanatomical architecture that regulates cortical excitability across distinct timescales. The first component is the brainstem ascending reticular system, including midbrain dopaminergic pathways (e.g., the tecto-nigral pathway from the superior colliculus), which convey rapid sensory signals and set a tonic, state-dependent excitability baseline for the motor network^1,2^. The second component consists of cortical top-down projections from the dorsal premotor cortex, supplementary motor area, and anterior cingulate cortex to the primary motor cortex^3,4^. These projections add a phasic, task-specific excitability boost during novel, complex, or effortful motor attempts, even in the absence of high mechanical load. This dual architecture implies that excessive GMA—and the resulting selectivity dilution— can arise from two convergent sources: mechanical necessity (via brainstem upregulation) and cognitive/attentional effort (via cortical top-down drive), a distinction with direct implications for both motor learning and rehabilitation. We use GMA as a phenomenological variable for changes in excitability that may involve several ascending and cortical pathways. The mapping from these pathways to eligible-pool size, and its dependence on mechanical or cognitive demands, remains a hypothesis requiring direct measurement. Furthermore, such GMA-regulated learning systems inherently tend to reinforce those loops capable of executing the corresponding commands or inputs earlier under lower GMA support; this, in turn, inevitably gives rise to competition among different loops and the switching between the effective loops under varying conditions.

### SI 2. Biological correspondence of the abstract network

In Figure 1A, the upstream population (Sb) represents neuronal ensembles in the primary somatosensory cortex (S1), whose axons project to Layer 2/3 of M1. The downstream output units (Ma to Md) correspond to the cascade of M1 Layer 5 corticospinal neurons and spinal α-motoneurons, which ultimately determine net muscular torque. During early development, this S1→M1 pathway constitutes a fundamental cortico-cortical reflex arc. With maturation and repeated successful performance, circuits involving higher-order cortical areas (e.g., premotor, supplementary motor, and posterior parietal cortices) gradually form new low-threshold pathways that functionally outcompete and progressively dominate over the reflexive S1→M1 loops, thereby shifting the system toward the Left Edge of the SRW.

S_b_ and M_a_–M_d_ are abstract input and output populations inspired by sensorimotor pathways. Each sampled element represents an effective output contribution. The developmental replacement of diffuse pathways by selectively recruited pathways is a proposed biological interpretation, not a result of the sampling model.

For extensor-targeted movements in a flexor-dominant system, there exists a narrow window of intermediate *n* where both conditions are satisfied: *P*_*success*_ > *θ* (sufficient success probability to trigger reinforcement, here set to 0.1, which means that on average, about one out of ten attempts will succeed: below it, successful trials may rare to drive reliable reinforcement within a behaviorally relevant time window) and *S*_*R*_(*n*) > *ϕ*_*sel*_ (sufficient selectivity for meaningful learning, here *ϕ*_*sel*_ set to 0.05). The window narrows with increasing *R* (Figure 1G red curve) and may disappear when *R* is too high (Figure 1J red curve and SI 3). The threshold *ϕ*_*sel*_ is system- and task-specific, set above the population baseline *p*_1_, demarcating the level above which reinforcement confers a detectable specificity advantage over random co-activation. Define *W*_*R*_(*θ*) = {*n* : *P*_*R*_(*n*) > *θ*} and *L*_*R*_(*θ, ϕ*_*sel*_) = {*n* : *P*_*R*_(*n*) > *θ* and *S*_*R*_(*n*) > *ϕ*_*sel*_}, with *θ* = 0.1 used illustratively and *ϕ*_*sel*_ = 0.05 specified for each analysis. No selective-window width is asserted without specifying *ϕ*_*sel*_. Extension results are shown in Figure 1H–J.

Crucially, the emergence of structured functional connectivity does not abolish the dilution constraint; it merely lowers the effective GMA level required to generate a given output torque. In a mature network, where task-relevant synapses have been selectively strengthened, a smaller co-activated ensemble suffices to produce *F*_*n*_ > *R*. Healthy individuals therefore operate naturally at the Left Edge (e.g., the low-*n* portion of the blue curves in Figure 1G, J), where selectivity is preserved and non-selective co-activation is weakened. We hypothesize that learning may reduce the GMA and eligible-pool size needed for successful performance. Structured recruitment changes the sampling distribution and therefore requires a separate model; operation near the selective region in healthy individuals remains to be tested.

This Left Edge strategy is actively enforced by a competitive refinement among low-threshold pathways—functionally pre-strengthened pathways that can be recruited with minimal GMA (see Glossary for full definition). Consequently, as GMA rises gradually from baseline, these low-threshold pathways with pre-strengthened connectivity are preferentially recruited; successful task completion terminates motor instruction with minimal delay, limiting the opportunity for higher-threshold pathways (that require higher GMA levels) to receive co-activation and reinforcement. The preferentially activated low-threshold pathways thus receive exclusive differential strengthening, which further enhances their synaptic efficacy and progressively lowers the GMA level required for their future recruitment. We hypothesize that recruitment order and rapid termination after success can favor earlier recruited pathways. The dynamic ensemble model implements success-gated reinforcement, failure-dependent depression, and competitive pruning, but it does not include explicit feedback delay, eligibility-trace decay, or recruitment order. Testing the proposed mechanism requires these additional dynamics.

Convergent M1 evidence is consistent with this competitive refinement, which we propose as a testable circuit-level hypothesis. The evidence includes: learning restructures cortical ensembles toward task-predictive states^5^; anti-correlated populations exert winner-take-all exclusion of higher-threshold pathways^6^; chandelier cells suppress task-irrelevant activity^7^; and dopamine-dependent reorganization of Layer 5b ensembles progressively enhances their movement encoding and lowers the drive required for their recruitment^8^. Collectively, these findings suggest that multiple complementary mechanisms may converge to implement the competitive refinement we describe, though direct demonstration of low-threshold pathways as discrete anatomical entities awaits further experimental validation. These studies motivate a hypothesis of competitive refinement, but do not by themselves identify the model’s low-threshold pathways or establish a reduction in GMA. Direct tests should measure recruitment thresholds, task-specific eligibility, and the timing of feedback within the same circuit.

Through repeated cycles, this self-reinforcing consolidation strongly favors a network configuration that responds to the same stimulus with progressively lower GMA and minimal energy expenditure. Yet this metabolic optimality comes at a cost: the systematic exclusion of alternative pathways renders the network exquisitely specialized but structurally brittle, with greatly diminished latent reserve capacity—a vulnerability that becomes catastrophic when the dominant low-threshold pathways are disrupted by lesions such as stroke. Repeated selective recruitment may reduce the drive required for performance. Whether this also reduces metabolic cost or reserve capacity is an additional hypothesis; these quantities are not represented in the present model.

### SI 3. Mathematical Derivations of Success Probability and Selectivity Dilution

This note provides the mathematical foundation for the selectivity dilution framework presented in the main text. We derive the exact finite-population expressions for two core quantities: (1) the probability of successful movement initiation, *P*_*success*_(*n*), and (2) the conditional proportion of task-relevant synapses in successful trials, *S*_*R*_(*n*)—the formal metric of selectivity dilution.

We also provide an analytical approximation based on the bivariate truncated normal distribution. This approximation is used only for intuitive understanding of the qualitative trend and is not used for the final numerical ranges. All reported *n* ranges were obtained by exact multivariate hypergeometric summation on a fine grid of *n* (e.g., *n* = 1,…,500 for *R* = 2, 8 and *n* = 1,…,5300 for *R* = 32). Monte Carlo simulations were used only for graphical validation. Evaluate every integer *n* = 1,…,5300. We distinguish numerical evaluation of exact combinatorial formulas from exact-integer verification.

#### 3.1. Model Setup and Notation

The synaptic pool consists of *N* = 5300 elements, each with a discrete force contribution. These values are illustrative, not fitted to a specific muscle or cortical population. The composition is: The values specify an illustrative parameter set. Monotonicity of *S*_*R*_(*n*) is a property to be verified for this population and resistance range.

| Type | Force (a.u.) | Count | Notation |
| --- | --- | --- | --- |
| Extensor (task-relevant) | +9 | 100 | Target class |
| Flexor (moderate) | −6 | 100 | Non-target |
| Flexor (strong) | −9 | 100 | Non-target |
| Task-irrelevant | 0 | 5000 | Non-target |

On each trial, a subset of *n* synapses is randomly co-activated without replacement—a reasonable approximation for nascent circuits prior to task-specific refinement. The net force generated is the sum of the sampled synaptic weights:

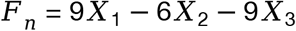

where *X*_1_: number of sampled +9 (extensor) synapses; *X*_2_: number of sampled −6 (moderate flexor) synapses; *X*_3_: number of sampled −9 (strong flexor) synapses. The trial is considered successful if *F*_*n*_ > *R*, where *R* is the external resistance threshold.

For each integer *n* ∈ {1,…,5300}, let *A*_*n*_ be a uniformly sampled n-element subset of the fixed population. Define *X*_*j,n*_ = |*A*_*n*_ ∩ *C*_*j*_| for the four classes *C*_*j*_, so

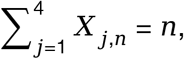

and set *F*_*n*_ = 9*X*_1,*n*_ − 6*X*_2,*n*_ − 9*X*_3,*n*_. The weights represent effective torque contributions in arbitrary units; *R* is expressed in the same units. Samples for different *n* need not be independent.

The varying torque values are also related to the differing force arms resulting from the different positions of the muscle tendon attachment points, in addition to being related to differences in motor unit size across different muscle groups. The size principle characteristic of motor unit activation also leads to a larger variance in output torque at high GMA levels; this, in turn, facilitates leveraging the fluctuations in this variance to increase the probability of *F*_*n*_ > *R*. The present model uses uniform recruitment and does not implement the size principle. Its torque variance is *nσ*^2^ (*N* −*n*)/(*N* − 1), which increases up to *n* = *N*/2 and decreases thereafter. Changes in success probability depend on the mean, variance, threshold, and full distribution, rather than variance alone.

#### 3.2. Exact Definitions (Multivariate Hypergeometric Model)

The joint distribution of *X*_1_, *X*_2_, *X*_3_ is multivariate hypergeometric:

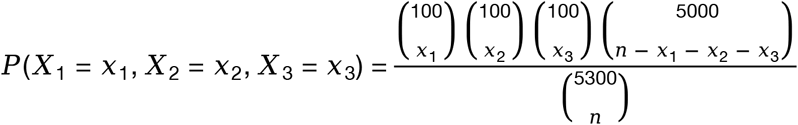

with the constraints:

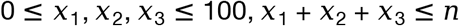

Let *D*_*n*_ = {(*x*_1_, *x*_2_, *x*_3_) ∈ *Z*^3^ : 0 ≤ *x*_*j*_ ≤ 100 for *j* = 1, 2, 3 and 0 ≤ *n* − *x*_1_ − *x*_2_ − *x*_3_ ≤ 5000}. Here *n* ∈ {1,…,5300}. All sums below are over *D*_*n*_; equivalently, define *C*(*m, k*) = 0 for noninteger *k* or *k* ∉ {0,…,*m*}. The number of sampled zero-force synapses is implicitly *x*_4_ = *n* − *x*_1_ − *x*_2_ − *x*_3_.

##### 3.2.1 Success Probability

The probability that a random co-activation of *n* neurons generates sufficient force to overcome resistance *R* is the cumulative sum over all combinations satisfying the force threshold:

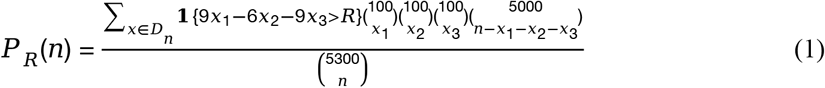

This is the exact expression. It must be evaluated numerically as a triple summation over the feasible domain. The computation is feasible for the relevant *n* range as *x*_1_, *x*_2_, *x*_3_ ≤ 100. Define *w*_*n*_(*x*) = *C*(100, *x*_1_)*C*(100, *x*_2_)*C*(100, *x*_3_)*C*(5000, *n* − *x*_1_ − *x*_2_ − *x*_3_). Then *P*_*R*_(*n*) = *Σ*_*x*∈*Dn*_ 1{9*x*_1_ − 6*x*_2_ − 9*x*_3_ > *R*} *w*_*n*_(*x*)/*C*(5300, *n*). The success event is strictly greater than *R*. Because force lies in 3*Z*, it is equivalent to *F*_*n*_ ≥ 3(⌊*R*/3⌋ + 1). The expression can be evaluated by direct summation or by grouping the three active classes according to *k* = *x*_1_ + *x*_2_ + *x*_3_ and then convolving with *C*(5000, *n* − *k*). The latter gives the same exact counts with less repeated computation.

##### 3.2.2 Selectivity Dilution Metric

The core metric of interest is the expected proportion of task-relevant (extensor) synapses among the co-activated set, conditional on successful force generation:

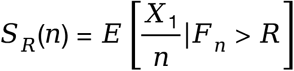

Expanding the conditional expectation yields the exact expression:

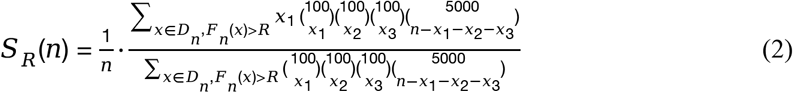

For *n* ≥ 1 with *P*_*R*_(*n*) > 0, define *S*_*R*_(*n*) = *E*[*X*_1,*n*_/*n* | *F*_*n*_ > *R*] = [*Σ*_*x*∈*Dn, Fn*(*x*)>*R*_ *x*_1_ *w*_*n*_(*x*)] / [*n Σ*_*x*∈*Dn, Fn*(*x*)>*R*_ *w*_*n*_(*x*)]. Set *S*_*R*_(*n*) undefined when *P*_*R*_(*n*) = 0. The SRW is the separate set *W*_*R*_ = {*n* : *P*_*R*_(*n*) > 0.1}. In general *S*_*R*_(*n*) is not *S*_*R*_(1)/*n* because *E*[*X*_1,*n*_ | *F*_*n*_ > *R*] also depends on *n*. Interpretation: *S*_*R*_(*n*) is not a heuristic or empirical curve; it is the mathematically exact conditionally expected composition of the reinforced ensemble. Numerical evaluation of this exact expression shows the monotonic decline of *S*_*R*_(*n*) as *n* increases, and its numerical approach toward the population baseline *p*_1_ within the feasible range where *P*_*success*_(*n*) remains non-negligible. Equation (2) is an exact definition. For the specified extension population, exhaustive exact-integer comparisons verify strict decrease for all real 2 ≤ *R* ≤ 12 on the positive-probability domain (see the audit addendum). This is a finite computer-assisted verification, not a parameter-free analytic theorem. Within the SRW, *S*_*R*_(*n*) can remain substantially above *p*_1_. For *R* = 2, *n* = 410, *S*_*R*_ ≈ 0.027293, while *p*_1_ ≈ 0.01887, still about 44.7% higher.

#### 3.3 Analytical Approximation (Bivariate Truncated Normal Framework)

The exact triple sum is the definitive numerical method. The following bivariate truncated normal approximation is provided only as an analytical tool to illustrate the qualitative behavior of *S*_*R*_(*n*). It is not used for the final numerical ranges. In the parameter regime where *P*_*success*_(*n*) is non-negligible, the joint distribution of (*X*_1_, *F*_*n*_) can be approximated by a bivariate normal distribution with finite population correction. The Gaussian approximation requires separate validation of the joint distribution and its truncated tail. A non-negligible success probability alone is insufficient. We use the exact discrete formulas for probabilities, conditional means, and window boundaries, and report approximation errors over any range where the Gaussian expression is interpreted quantitatively. Impossible events and *n* = *N* are excluded from this approximation.

##### 3.3.1 Population Moments

Mean force per synapse:

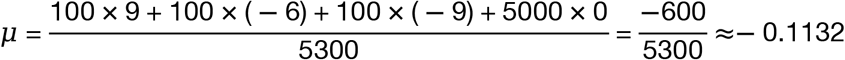

Variance of force per synapse:

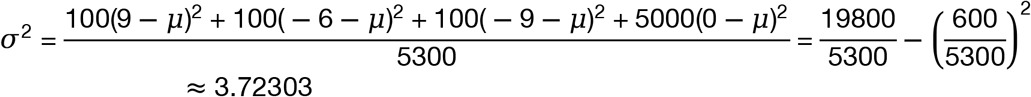

Proportion of extensor synapses:

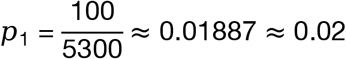

Here *σ*^2^ ≈ 3.72303 (population variance with divisor *N*).

##### 3.3.2 Sampling Moments (with Finite Population Correction)

For a sample of size *n* drawn without replacement, the marginal moments are: Mean of total force:

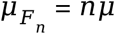

Variance of total force:

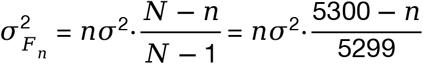

Mean of extensor count:

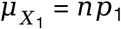

Covariance between extensor count and total force:

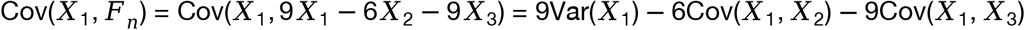

Using the standard variance and covariance formulas for the multivariate hypergeometric distribution,

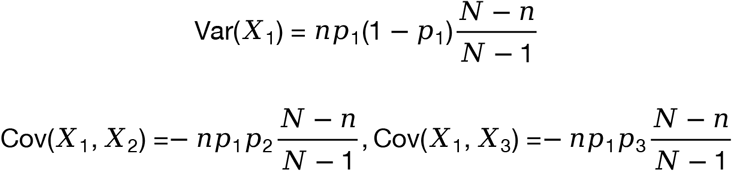

Substituting these into the expression above yields

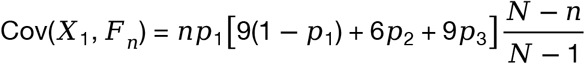

Since *μ* = 9*p*_1_ − 6*p*_2_ − 9*p*_3_, the bracket simplifies to 9 − *μ*. The equality of the three class proportions is needed only for the numerical parameter values. Therefore,

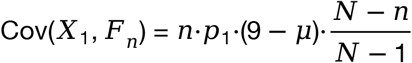

##### 3.3.3 The Conditional Expectation Formula

Using the properties of the truncated bivariate normal distribution (Tallis, 1961)^9^, the expected value of *X*_1_ given that *F*_*n*_ > *R* is:

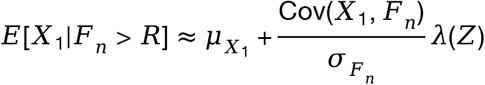

For a jointly Gaussian surrogate 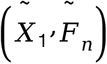 with the stated means and covariances,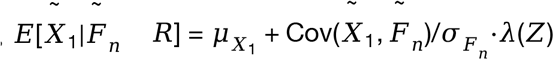. For the discrete sampling model, 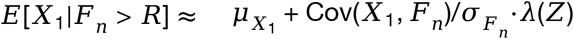. The finite-population correction matches moments; it does not make the discrete law Gaussian. In the above,

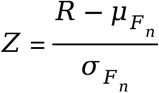

is the standardized threshold, and:

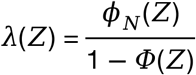

is the Inverse Mills Ratio^10,11^, with *ϕ*_*N*_ and *Φ* denoting the standard normal probability density function (PDF) and cumulative distribution function (CDF), respectively. The Inverse Mills Ratio captures the conditional intensity of the threshold exceedance; it is positive and strictly increasing with *Z*, since

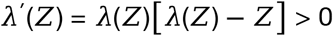

For a standard normal *G, λ(z*) =*E*[*G*|*G* >*z*] >*z*. Differentiating *λ*(*z*) = *ϕ*_*N*_(*z*)/(1 − *ϕ*(*z*)) gives *λ′*(*z*)= *λ(z)*[*λ(z)*−*z*]> 0. This derivative does not determine the sign of *dS*_*R*_/*dn* because both *Z*(*n*) and its prefactor depend on *n*. Use *ϕ*_*N*_ for the normal density and a distinct symbol *ϕ*_*sel*_ for the selectivity threshold. Dividing by *n* yields the analytical approximation for the dilution metric:

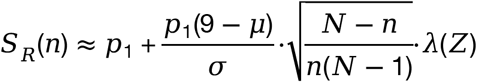

This expression is evaluable in a single line of code without nested summation. A technical note on its applicability: the conditional expectation formula above is a direct application of Tallis (1961) to the bivariate case. The inclusion of the finite population correction factor (*N*−*n*)/(*N*−1) in *σ*^2^_*Fn*_ is an adjustment for sampling without replacement from a finite synaptic pool. This adjustment is standard in finite-population sampling theory. Evaluate *λ* using a stable survival-function or log-survival-function implementation. Check the approximation against exact combinatorial results and report any violation as approximation failure.

#### 3.4. What the Analytical Form Reveals

The closed-form approximation above is not merely a computational convenience—it provides an intuitive explanation of the two central claims of the dilution framework.

##### 3.4.1 Monotonic Decay to Baseline

As *n* increases, the finite-population correction factor (*N*−*n*)/(*N*−1) eventually reduces *σ*^2^_*Fn*_ as *n* approaches *N*. Within the SRW, however, *σ*^2^_*Fn*_ typically still increases with *n*; the shrinkage only becomes relevant near the upper limit of the population. Within the feasible range where *P*_*success*_(*n*) remains non-negligible, numerical evaluation of the exact hypergeometric expression shows that *S*_*R*_(*n*) declines monotonically toward the population baseline *p*_1_. The limit *n* → *N* is not used as a formal conditional-expectation statement because *P*_*success*_(*n*) may vanish there. For this extension model, Var(*F*_*n*_) increases on *n* < 2650 and decreases on *n* > 2650. It increases throughout the reported SRWs. At *n* = 5300, *F*_*n*_ = −600 deterministically, so *S*_*R*_(5300) is undefined for *R* > 0. For fixed finite *N*, no conditional limit *n* → *N* is asserted. Within the *R* = 2 SRW, *S*_*R*_(410) ≈ 0.02729 remains above *p*_1_ ≈ 0.01887.

Numerical evaluation of the exact hypergeometric expression confirms that selectivity dilution is mathematically inescapable in this finite-population sampling system with a hard success threshold. The decline is not an artifact of simulation parameters or random seed. The observed decline is independent of Monte Carlo seeds because it is verified from combinatorial counts. It is established for the stated finite population and tested resistance values; robustness to changes in class sizes, weights, or recruitment rules requires additional analysis.

##### 3.4.2 The Narrow Selective Window

The formula also reveals why the selective window (where *S*_*R*_(*n*) > *ϕ*_*sel*_, with *ϕ*_*sel*_ denoting the minimum acceptable level of synaptic specificity for meaningful learning) is narrow and why it depends critically on *R*: the selectivity threshold *ϕ*_*sel*_ = 0.05 is an illustrative operational criterion rather than a derived biological requirement. Report its value, the resulting integer interval *L*_*R*_, and sensitivity to several values above the direction-specific baseline.

1. At low *n, σ*^2^_*Fn*_ is small, but *λ*(*Z*) is large because success is infrequent and requires extreme sampling fluctuations—these infrequent successes are dominated by extensor-rich samples, yielding high selectivity. At small *n*, successful configurations often require a high target fraction. This enrichment is quantified by the exact conditional count distribution. Its explanation need not rely on a Gaussian tail approximation, which can be inaccurate when the expected class counts are small.
2. At high *n*, within the feasible range where *P*_*success*_(*n*) remains non-negligible, success becomes increasingly compatible with samples close to the population average, so *S*_*R*_(*n*) approaches *p*_1_. In the extension model, if *n* becomes too large, the success event itself may vanish, and *S*_*R*_(*n*) is then undefined. In the extension model, larger *n* does not make success progressively easier: the negative population mean eventually dominates and success becomes rare or impossible. The decline of *S*_*R*_(*n*) must therefore be established separately from the success-probability trend. Proximity to *p*_1_ is reported quantitatively, not as a formal limit.

Increasing *R* shifts the threshold rightward and generally requires larger *n*, narrowing the selective window and forcing the system to operate at higher *n*, where dilution is more severe. For *R*_2_ > *R*_1_, {*F*_*n*_ > *R*_2_} ⊆ {*F*_*n*_ > *R*_1_}, hence 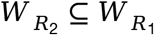. No corresponding inclusion follows for *L*_*R*_ because *S*_*R*_(*n*) also changes with *R*. In the discrete model, force is quantized in steps of 3, so small *R* changes can produce non-monotonic local shifts. For example, with *ϕ*_*sel*_ = 0.05, extension *L*_*R*_ is 7– 28 for *R* = 2 and 7–31 for *R* = 8. Across a wider range, increasing *R* generally shrinks *W*_*R*_ and eventually *L*_*R*_, but the selective region must be computed for each *R*; *R* = 10 cannot be inferred from *R* = 2 and *R* = 8 alone.

Nevertheless, exact numerical evaluation of the multivariate hypergeometric conditional expectation shows that *S*_*R*_(*n*) decreases monotonically within the SRW. A heuristic analytical argument, obtained by approximating the multivariate hypergeometric distribution with a multinomial distribution (valid for *n* ≪ *N*) and then applying the bivariate normal approximation, suggests the same trend; this argument is illustrative rather than a formal proof, and the exact monotonicity is confirmed numerically. For fixed *N* = 5300, class sizes (100, 100, 100, 5000), and weights (9, −6, −9, 0), strict decrease for every real *R* ∈ [2, 12] can be certified by exact-integer comparisons over all feasible *n*. The force lattice reduces this continuum of *R* to five distinct success events. Details and reproducible comparison formulas are given in the audit addendum; no general analytic monotonicity theorem is claimed.

#### 3.5. Parameters and Numerical Evaluation

For the specific network modeled in Figure 1 (*N* = 5300, 100 extensors (+9), 100 flexors (−6), 100 flexors (−9), 5000 null (0) synapses), the per-synapse moments are:

| Parameter | Value |
| --- | --- |
| $N$ | 5300 |
| $p_1$ | $100/5300 \approx 0.01887$ |
| $\mu$ | $-600/5300 \approx -0.1132$ |
| $\sigma^2$ | $\approx 3.72303$ |

Here *σ*^2^ = 3.72303 (population variance with divisor *N*). All numerical ranges below are computed from the exact multivariate hypergeometric expressions, Eqs. (1) and (2), with the success probability threshold set to 0.1. The resulting integer *n* ranges are:

| Resistance $R$ | Success event | Integer $n$ range |
| --- | --- | --- |
| 2 | $F > 2$ | $7 \leq n \leq 410$ |
| 8 | $F > 8$ | $7 \leq n \leq 299$ |
| 32 | $F > 32$ | No solution |

Boundary checks: *R* = 2 gives *P*(6) ≈ 0.09816, *P*(7) ≈ 0.11133, *P*(410) ≈ 0.10017, *P*(411) ≈ 0.09989. *R* = 8 gives *P*(6) ≈ 0.08966, *P*(7) ≈ 0.10006, *P*(299) ≈ 0.10005, *P*(300) ≈ 0.09979. The inequality is strictly *P* > 0.1. For *R* = 32 (high resistance), the exact calculation confirms that *P*_*success*_(*n*) ≤ 0.1 for all *n* ∈ {1,…,5300}. The maximum success probability is below 0.1, so no sustainable reinforcement window exists under this condition. For *R* = 32, max_*n*_*P*_*R*_(*n*)≈ 0.02192 at *n* = 253, so *W*_*R*_(0.1) is empty. Success remains possible, and *S*_*R*_(*n*) is defined on *n* = 4,…,5229. The choice *θ* = 0.1 corresponds to an expected 10 attempts to first success under independent trials. For many animal learning paradigms, a success rate below 0.1 would require more than ten trials per reinforcement under independent sampling, which may exceed the behaviorally relevant time window for early motor learning. This motivates *θ* = 0.1 as an illustrative cutoff, not a derived biological constant. Sensitivity to *θ* = 0.05, 0.1, 0.2 should be reported.

These exact numerical results directly support the breadth-precision trade-off illustrated in Figure 1 and discussed in the main text. These results quantify success opportunities in the specified static model. The implications for synaptic learning dynamics and rehabilitation are hypotheses that require explicit update rules and empirical testing.

### SI 4. Monotonicity of SR(n) in Flexion and Extension Training

SI4 applies the exact definitions of SI3 to the two movement directions separately. In our framework, the task-relevant synapse population is defined by its contribution to the target movement direction:

1. Extension training: task-relevant synapses are those from triceps motor units, each contributing +9 to the extension torque. Flexor synapses (biceps, −9; brachioradialis, −6) and task-irrelevant synapses (0) are non-target.
2. Flexion training: task-relevant synapses are those from biceps (+9) and brachioradialis (+6) motor units. Extensor synapses (triceps, −9) and task-irrelevant synapses (0) are non-target.

For extension training, the task-relevant synapses are defined as those that contribute positively to the target torque, while non-target synapses contribute non-positively. The success condition is *F*_*n*_ > *R*, where *R* > 0 is a positive resistance threshold in the target direction.

Under this unified definition, numerical evaluation of the exact conditional expectation shows that the conditional proportion of task-relevant synapses on successful trials, *S*_*R*_(*n*) = *E*[*r*_*n*_/*n* | *F*_*n*_ > *R*], is monotonically decreasing with *n* within the SRW. The bivariate truncated normal approximation provides an intuitive explanation for this trend, but the exact monotonicity is established numerically from the multivariate hypergeometric distribution. The two directions are evaluated separately. In the specified model, extension selectivity is strictly decreasing on its feasible domain for the resistance ranges verified below. Flexion selectivity is non-increasing for *R* = 2, 8, 12, 22, 32 with a terminal plateau at 200/5300 when every possible sample succeeds.

#### 4.1.Corrected Conditional Expectation

Using the truncated bivariate normal approximation, the conditional expectation is

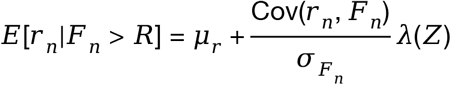

For the discrete model,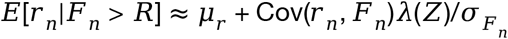. Equality holds only for a jointly Gaussian surrogate with these moments. In the above,

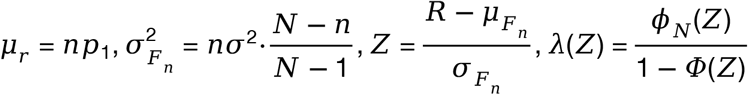

and

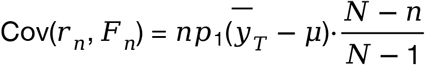

with 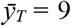 for extension training and 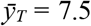 for flexion training (biceps +9 and brachioradialis +6). Here *μ* denotes the population mean force in the target direction. For extension, *μ* = −600/5300 ≈ −0.1132; for flexion, the sign of the target direction is reversed, so *μ* must be recomputed accordingly. For extension, *r*_*n*_ = *X*_1,*n*_, *F*_*n*_ = 9*X*_1,*n*_ − 6*X*_2,*n*_ − 9*X*_3,*n*_, *p*_*T*_ = 100/5300, *μ* = −600/5300, and 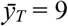. For flexion, *r*_*n*_ = *X*_2,*n*_ + *X*_3,*n*_, *F*_*n*_ = −9*X*_1,*n*_ + 6*X*_2,*n*_ + 9*X*_3,*n*_, *p*_*T*_ = 200/5300, *μ* = +600/5300, and 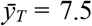. Both directions have *σ*^2^ ≈ 3.72303. In either direction 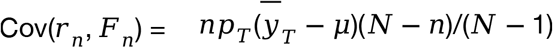. Therefore

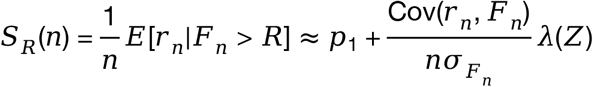

For extension training this becomes

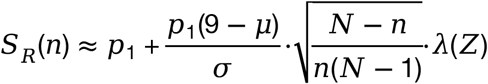

The exact monotonicity is established numerically from the exact hypergeometric conditional expectation. Report the resistance set, positive-probability domain, and all adjacent comparisons. Use integer cross-multiplication of the exact count sums to distinguish strict decrease from equality; do not decide monotonicity by rounded plotted values.

#### 4.2 Numerical Verification of Monotonicity

Exact evaluation of the conditional expectation in Eq. (2) for a wide range of *n* within the SRW shows that *S*_*R*_(*n*) decreases monotonically as *n* increases, and approaches the population baseline *p*_1_ within the feasible range where the success event has non-negligible probability. This monotonicity holds for both flexion and extension training, because in both cases task-relevant synapses are defined as those contributing positively to the target torque direction, ensuring Cov(*r*_*n*_,*F*_*n*_) > 0. The only difference lies in which synaptic populations are labeled as “task-relevant,” which affects the numerical values of *p*_1_ and the parameters in the approximation, but not the sign of the monotonic trend. In our specific network, *p*_1_ = 200/5300 ≈ 0.04 for flexion training (biceps + brachioradialis) and *p*_1_ = 100/5300 ≈ 0.02 for extension training (triceps only), which explains why the two curves in Figure 1 converge to different baselines. The two directions are evaluated separately. In the specified model, extension selectivity is strictly decreasing on its feasible domain for the resistance ranges verified below. Flexion selectivity is non-increasing for *R* = 2, 8, 12, 22, 32 with a terminal plateau at 200/5300 when every possible sample succeeds.

Positive Cov(*r*_*n*_,*F*_*n*_) does not prove monotonicity in *n*. For flexion, replace the event in Eq. (2) by −9*x*_1_ + 6*x*_2_ + 9*x*_3_ > *R* and its numerator factor by *x*_2_ + *x*_3_. Exact-integer checks give non-increasing *S*_*R*_(*n*) for *R* = 2, 8, 12, 22, 32; once all samples succeed, *S*_*R*_(*n*) = 200/5300 exactly. For extension, the terminal success event instead disappears. These are distinct endpoint behaviors.

#### 4.3. Intuitive Interpretation

The Inverse Mills Ratio *λ*(*Z*)=*ϕ*_*N*_(*Z*)/(1−*ϕ(Z))*is strictly increasing in *Z* (Sampford, 1953; Mills, 1927)10,11, since

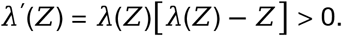

As *n* increases, 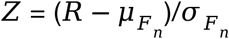 generally shifts rightward because 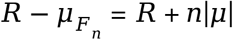 grows linearly while 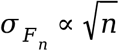 grows more slowly. Thus *λ*(*Z*) increases. At the same time, the sampling-fluctuation factor 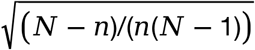 decreases with *n*. These two factors move in opposite directions, so their product is not guaranteed to be monotonic by this argument alone.

Nevertheless, exact numerical evaluation of the multivariate hypergeometric conditional expectation shows that *S*_*R*_(*n*) decreases monotonically within the SRW. The intuitive reason is that as *n* grows, success becomes increasingly compatible with samples close to the population average, so the conditional composition of successful samples approaches *p*_1_. The truncated normal approximation is only for intuition; the exact monotonicity is established numerically.

For extension, with *a* = 600/5300 and 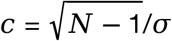,

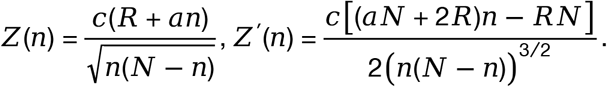

Thus *Z* decreases for *n* < *NR*/(*aN* + 2*R*) and increases thereafter; the turning point is 5300*R*/(600 + 2*R*), equal to 17.5497 for *R* = 2 and 68.8312 for *R* = 8. For flexion, 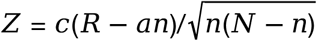 and the derivative numerator is (2*R* − *aN*)*n* − *RN*, which is negative for 2 ≤ *R* ≤ 32. Neither ^′^ > 0 nor the opposite movements of the two factors alone proves the exact discrete monotonicity.

### SI 5. Dynamic Ensemble-Based Structural Plasticity Model

SI5 introduces a separate dynamic model used for Figure 2; it does not replace the static results of SI3–4. The dynamic simulations in main Figure 2 and Figure SI1 use a separate ensemble-based structural plasticity model. This model is illustrative and is not fitted to experimental data. It is not the same as the static multivariate hypergeometric model in SI Sections 3–4.

#### Pools

For extension training, the total pool contains *N* = 5300 elements: 100 extensor elements with force +9, 100 biceps elements with force −9, 100 brachioradialis elements with force −6, and 5000 non-elbow elements with force 0. For flexion training, the signs are reversed: biceps +9, brachioradialis +6, triceps −9, non-elbow 0.

#### Reinforced pool

The reinforced pool *R*_*t*_ is a multiset of size 100. It is initialized by random sampling from the total pool. The non-reinforced pool *N*_*t*_ contains the remaining elements.

#### Activation

At trial *t*, GMA *G*_*t*_ determines the activated ensemble *A*_*t*_. The specific component is

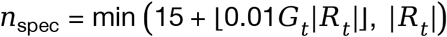

and the non-specific component is

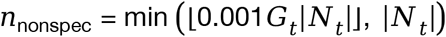

Elements are sampled randomly without replacement from *R*_*t*_ and *N*_*t*_, respectively. The activated force is

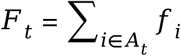

Success is *F*_*t*_ > *R*. **Update rules**. On success:

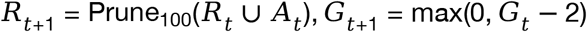

On failure:

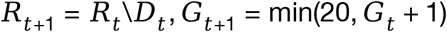

where *D*_*t*_ is a random subset of *A*_*t*_ ∩ *R*_*t*_ with removal rate

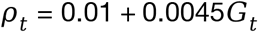

The modified condition in Figure 2G–J enforces *G*_*t*+1_ = max(*G*_*t*_ − 2,10) after trial 400 on successful trials. A run is labeled stabilized after 10 consecutive successful trials.

#### Simulation details

Each condition uses 20 independent runs, 500 trials per run, random seed 32. Solid lines are smoothed means using a moving average with window 9; shaded regions are 95% confidence intervals across runs; thin traces show the first five individual runs. The dotted line at GMA = 10 in baseline panels is a reference line and is not an enforced bound.

#### Limitations

This dynamic model does not include explicit synaptic weights, eligibility-trace decay, feedback delay, anatomical topography, motor-unit size principle, sensory feedback, spasticity mechanisms, or clinical dose–response. It therefore provides illustrative in silico support for hypotheses, not clinical proof. The dynamic model reports the composition of the reinforced pool over trials, not the static success-conditioned target fraction *S*_*R*_(*n*). Dynamic enrichment or bias should not be interpreted as a direct estimate of *S*_*R*_(*n*).

### SI 6. Summary

The exact formulas describe static uniform sampling with fixed contributions and no learning updates. The Gaussian formulas are heuristic approximations. Dynamic simulations in main Figure 2 and Figure SI 1 use a separate ensemble-based structural plasticity model specified in SI Section 5; they require the code, initialization, activation rule, reinforcement, depression, pruning, force generation, GMA control, stopping rules, and randomization described there. Ordered motor-unit recruitment is not implemented in either model. The dynamic model supports the plausibility of recruitment-dependent bias over trials, but it does not estimate *S*_*R*_(*n*) or establish clinical causality.

## Figures of SI

**Figure SI1.**
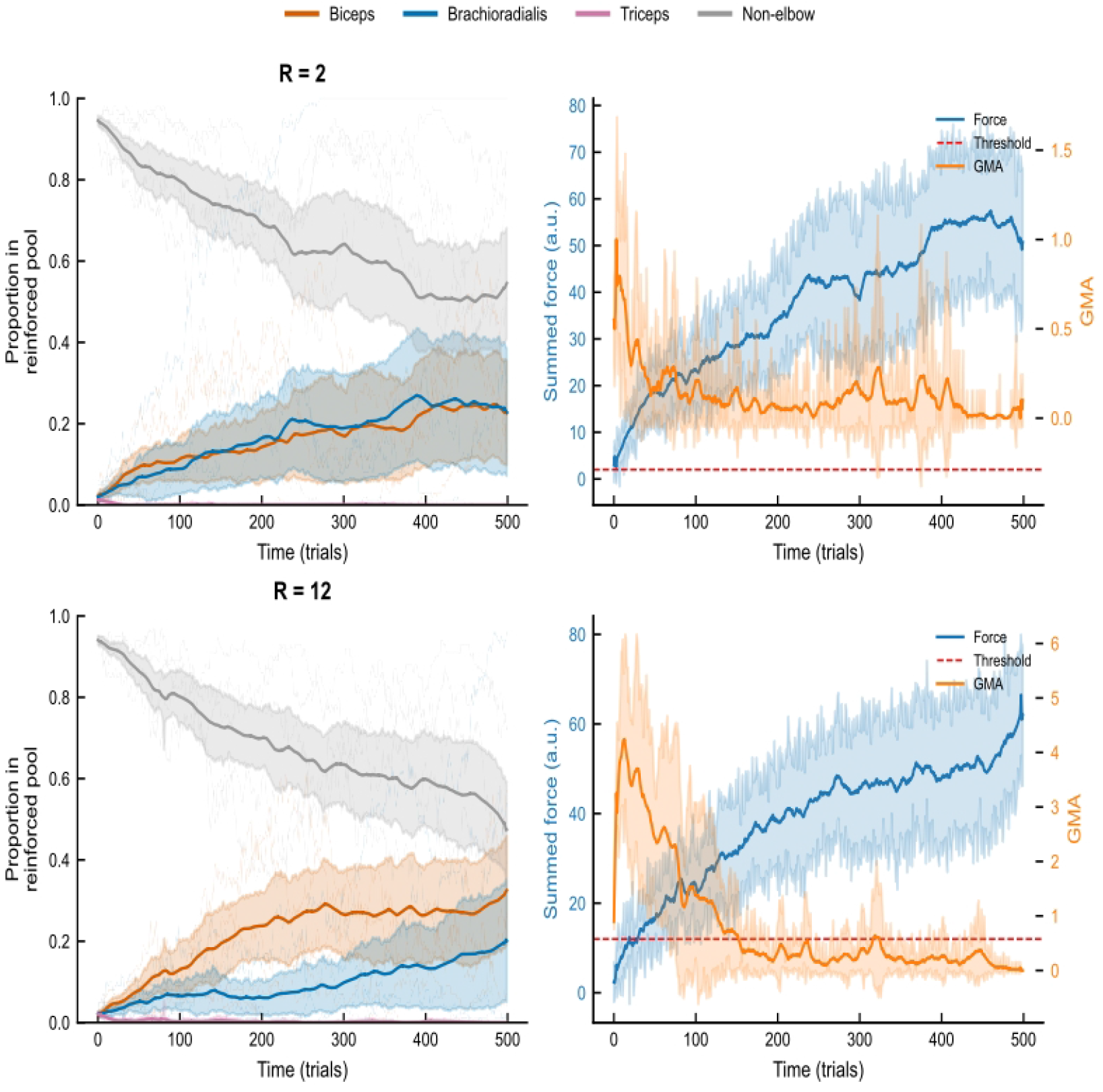
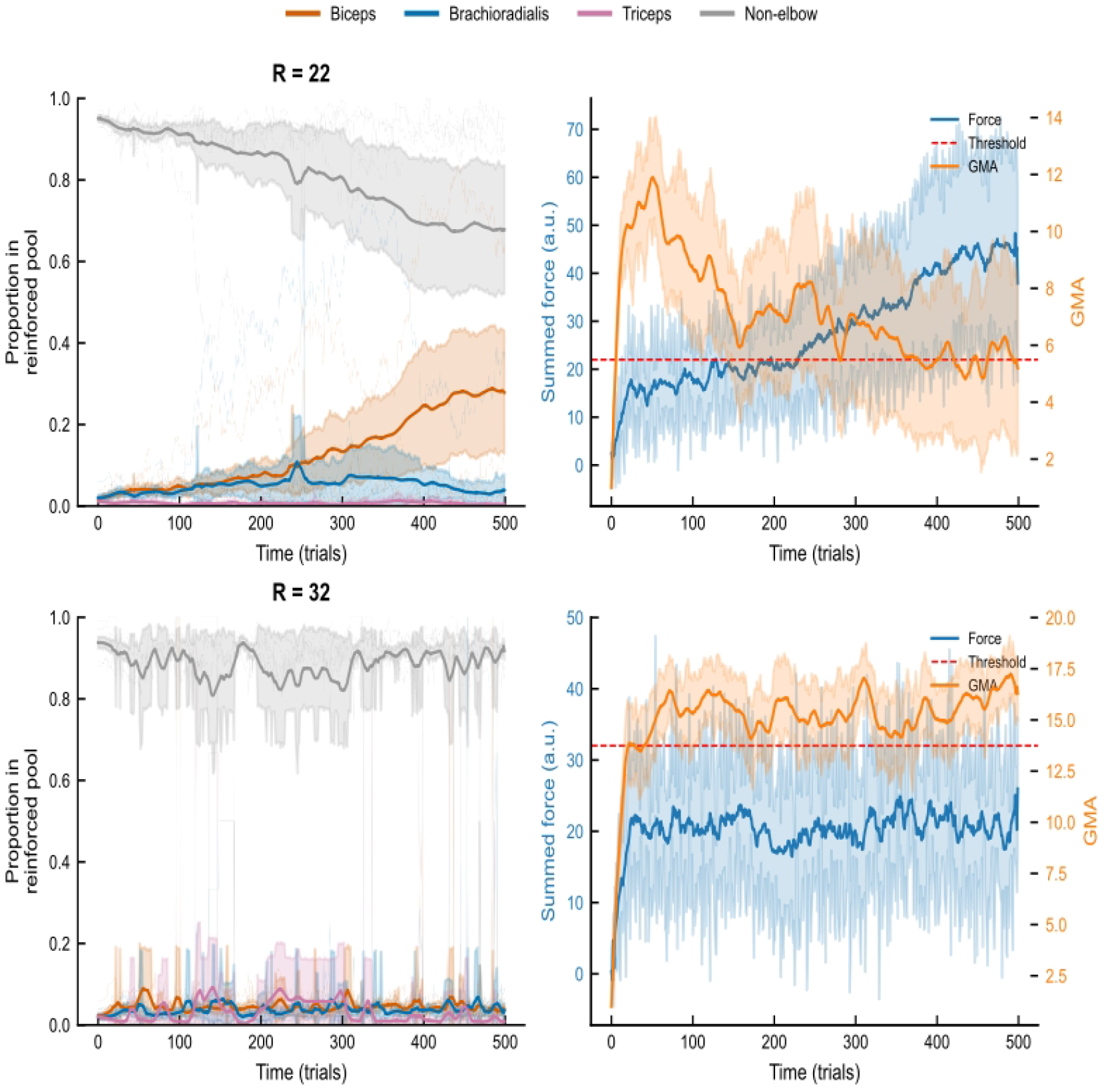
Competitive dynamics of synaptic strengthening under different resistance conditions in a flexor-biased network. Time course of the proportion of target muscles within the synaptic connections between Sb neurons and their downstream target muscles. The level of *R* that can be tolerated while strengthening the flexor muscles is higher (*R* = 2, 12, or 22 are all feasible, but *R* = 32 fails). Figure SI 1 illustrates outcomes of a separate dynamic learning model, whose full update rules, parameters, seeds, replicate count, and uncertainty-band definition are given in SI Section 5. Under the static flexion model, SRWs for *R* = 2, 12, 22, 32 are respectively 3–5300, 20–5300, 48–5300, and 83–5300. Failure to strengthen at *R* = 32 in the displayed dynamics must not be described as absence of a static success window. Report endpoint variability and sensitivity to the controller and learning rules.

**Figure SI2.**
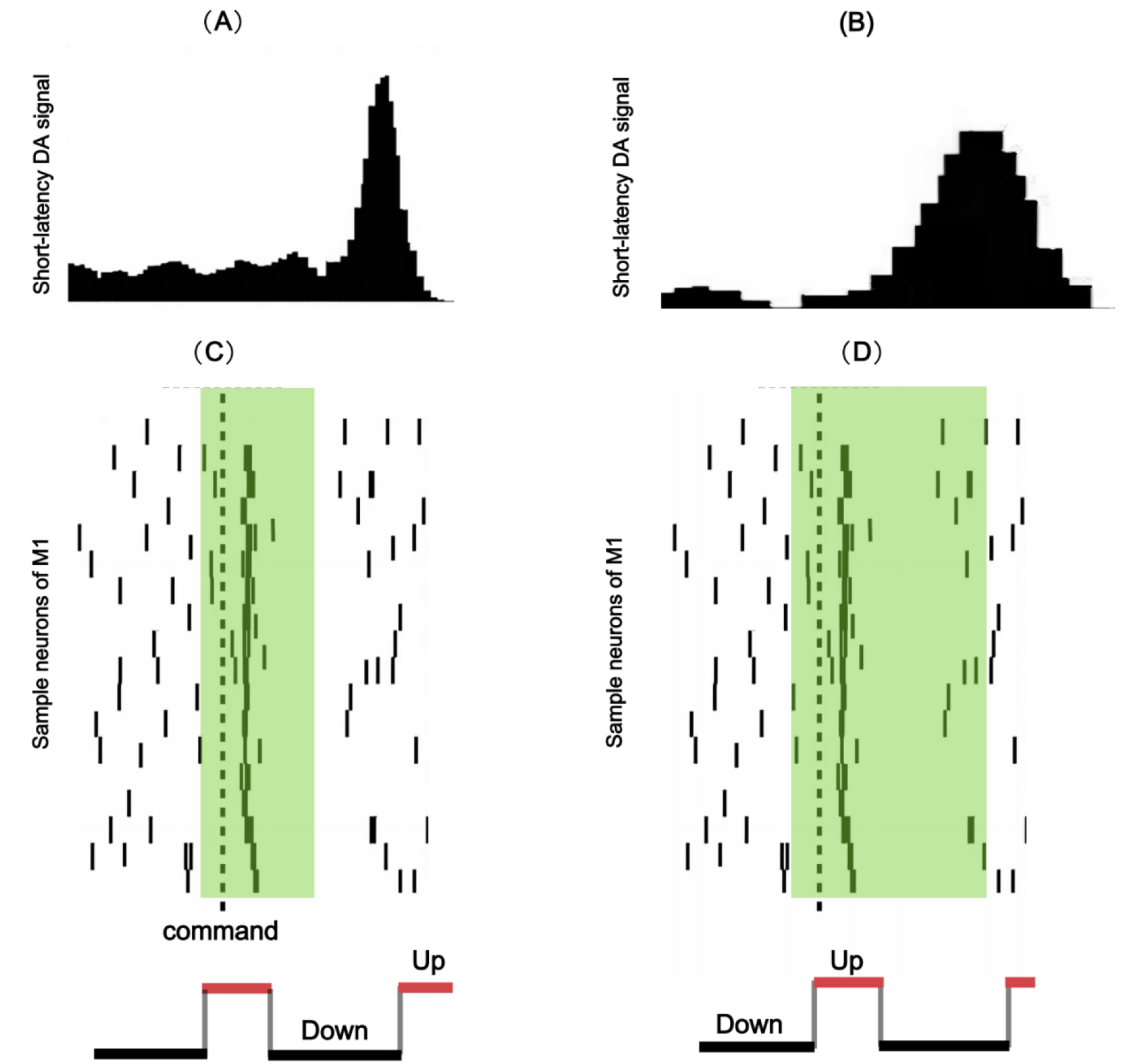
The phasic release pattern of dopamine and the potential impact of neural oscillations on selective dilution. The dopamine release peaks are steeper (A, C), and their corresponding enhancement time windows (green rectangle) are narrower (compared to B, D), which theoretically increases the *r*/*n* ratio. Neuronal oscillations concentrate neuronal firing activity within up-states, thereby helping to increase *n* within a given time window without substantially increasing overall brain energy consumption (C, D). Simultaneously, the subsequent down-states also reduce the likelihood of generating eligibility traces with irrelevant neurons to some extent. Figure SI 2 is a conceptual schematic. A narrower modulatory window may improve selectivity only if target-related eligibility is temporally more concentrated than non-target eligibility. Oscillations may change recruitment and eligibility timing, but their net effect on *S*_*R*_(*n*) depends on these distributions. Energetic consequences are not modeled here. Label the trace amplitudes and time axes as schematic arbitrary units.

**Figure SI3.**
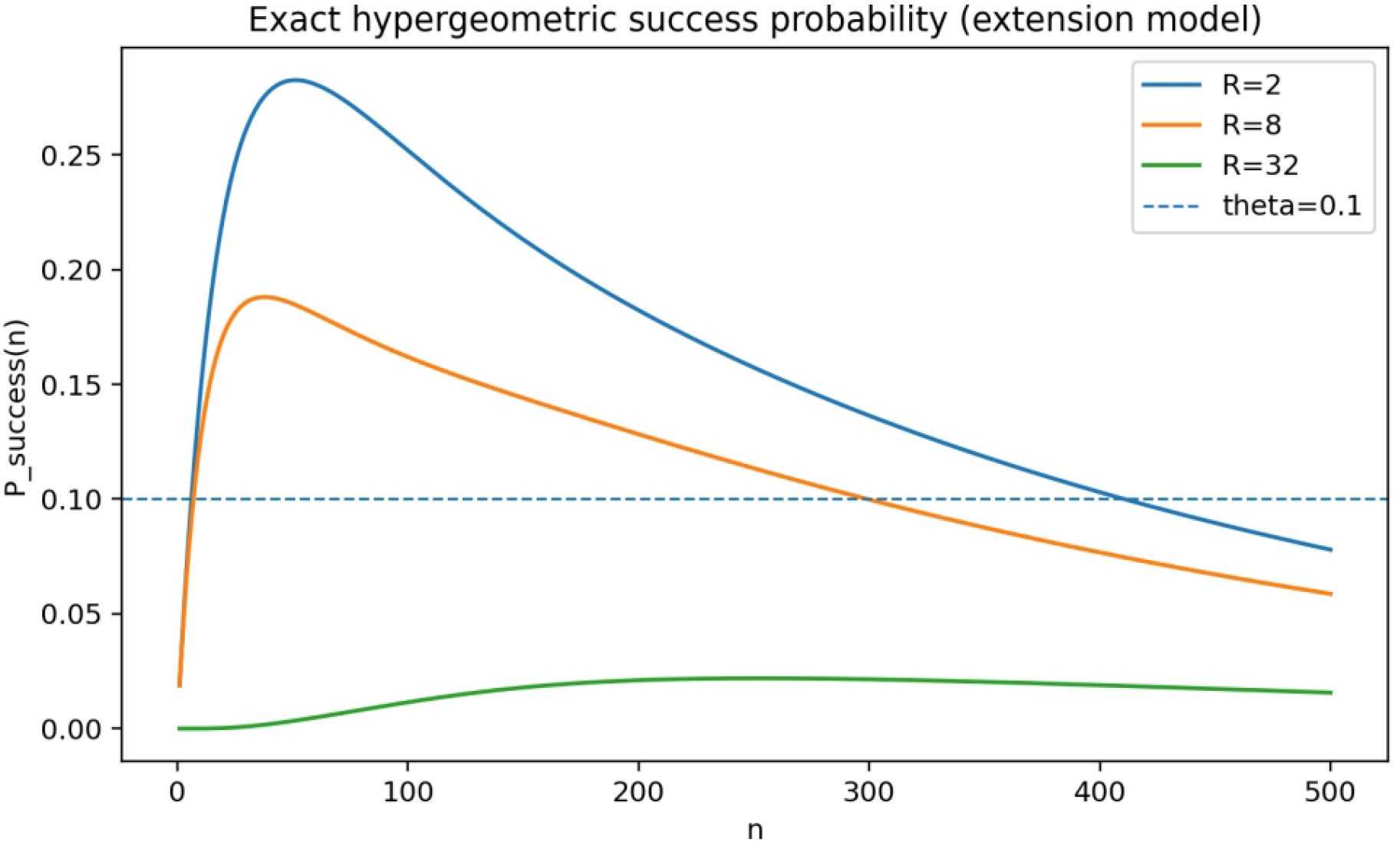
**Curves illustrating the variation of the extension success probability with increasing *n* under different R levels, calculated using the multivariate hypergeometric model**. The dashed line corresponds to θ = 0.1.

### Audit addendum

#### Exact counting method

*C*(*m, k*) = 0 if *k* ∉ {0,…,*m*}. For *a, b, c* = 0,…,100, set *k* = *a* + *b* + *c, f* = 9*a* − 6*b* − 9*c, w* = *C*(100, *a*)*C*(100, *b*)*C*(100, *c*). For extension, event *f* > *R*, target count *a*. For flexion, event *f* > *R*, target count *b* + *c*. Accumulate *A*_*R*_(*k*) and *B*_*R*_(*k*). For each *n* = 1,…,5300, compute *T*_*R*_(*n*) = *Σ*_*k*=0..300_ *A*_*R*_(*k*)*C*(5000, *n* − *k*), *U*_*R*_(*n*) = *Σ*_*k*=0..300_ *B*_*R*_(*k*)*C*(5000, *n* − *k*). Then *P*_*R*_(*n*) = *T*_*R*_(*n*)/*C*(5300, *n*), *S*_*R*_(*n*) = *U*_*R*_(*n*)/(*n T*_*R*_(*n*)), defined only when *T*_*R*_(*n*) > 0. All intermediate quantities are non-negative integers

#### Exact comparisons

*P*_*R*_(*n*) > 0.1 is equivalent to 10*T*_*R*_(*n*) > *C*(5300, *n*). For adjacent *n* with *T* > 0, *S*_*R*_(*n*+1) < *S*_*R*_(*n*) is equivalent to *D*_*R*_(*n*) = (*n*+1)*T*_*R*_(*n*+1)*U*_*R*_(*n*) − *nT*_*R*_(*n*)*U*_*R*_(*n*+1) > 0; equality is *D* = With *ϕ*_*sel*_ = 1/20, *S*_*R*_(*n*) > *ϕ*_*sel*_ is equivalent to 20*U*_*R*_(*n*) > *nT*_*R*_(*n*). Use arbitrary-precision integers; no floating-point comparison is needed.

#### Extension certification for 2 ≤ R ≤ 12

Because *F*_*n*_ ∈ 3*Z, R* ∈ [2, 3], [3, 6], [6, 9], [9, 12], {12} correspond to *F*_*n*_ ≥ 3, 6, 9, 12, 15. Representative *R* = 2, 3, 6, 9, 12 cover the whole interval. Positive-probability domains are 1–5233, 1–5232, 1–5232, 2–5232, 2–5231. For all 26,153 adjacent comparisons, integer *D* is positive. This is a finite computer-assisted proof for fixed parameters, not an analytic theorem for arbitrary populations. The corresponding extension SRWs for *θ* = 0.1 are 7–410, 7–357, 7–299, 50–228, empty. *R* = 22, 32 extension SRWs are also empty; max *P* ≈ 0.044944287 at *n* = 193 and 0.021923362 at *n* = 253, respectively. *R* = 12 has max *P* = 0.093041419 at *n* = 135. These decimals are rounded numerical results; integer comparisons and decimal reports must be stated separately.

#### Flexion full check

For *R* = 2, 8, 12, 22, 32, checking all *n* = 1,…,5300 shows no increase; SRWs are 3–5300, 6–5300, 20–5300, 48–5300, 83–5300. Strict decrease ends at *n* = 5234, 5235, 5235, 5236, 5237, after which *S*_*R*_ is constant at 200/5300. Thus the global statement is non-increasing, not strictly decreasing.

#### Plateau explanation

For flexion, total force over all elements is 600. If a sample omits *m* = *N* – *n* ≤ 100 elements, the maximum omitted force is 9*m*, so the minimum possible *F*_*n*_ = 600 − 9*m*. When 600 − 9*m* > *R*, every sample succeeds, *P* = 1, and *S*_*R*_ = *E*[*r*_*n*_]/*n* = 200/5300. This gives the plateau onset and shows that Cov > 0 does not imply global strict decrease.

#### Selective-window counterexamples

With *θ* = 0.1, *ϕ*_*sel*_ = 0.05, extension selective windows are 7–28 for *R* = 2 and 7–31 for *R* = 8; flexion windows are 3–55, 6–77, 20–102, 48–143, 83–180 for *R* = 2, 8, 12, 22, 32. These are not clinical thresholds; they refute the general claim that increasing *R* necessarily shrinks the selective window.

#### Dynamic method checklist

At minimum: Sb and output pool sizes; connection initialization distribution; eligibility-trace generation, sign, amplitude, decay; DA/success timing; weight updates on success and failure; forgetting, competition, normalization; mapping from weights to actual torque; GMA control equation and mapping to *n*; timing of imposed lower bounds; random seeds; replicate counts; shading and smoothing definitions; raw arrays for each panel. Until these are supplied, dynamic panels cannot be independently certified, and static *S*_*R*_ cannot be used to infer dynamic steady states.

## References

1. Reinoso-Suárez, F., De Andrés, I. & Garzón, M. Functional Anatomy of the Sleep-Wakefulness Cycle: Wakefulness. vol. 208 (Springer Berlin Heidelberg, Berlin, Heidelberg, 2011).

2. Groppa, S. et al. The human dorsal premotor cortex facilitates the excitability of ipsilateral primary motor cortex via a short latency cortico-cortical route. Hum. Brain Mapp. 33, 419–430 (2012).

3. Khachouf, O. T., Chen, G., Duzzi, D., Porro, C. A. & Pagnoni, G. Voluntary modulation of mental effort investment: an fMRI study. Sci. Rep. 7, 17191 (2017).

4. Munn, B. R., Müller, E. J., Wainstein, G. & Shine, J. M. The ascending arousal system shapes neural dynamics to mediate awareness of cognitive states. Nat. Commun. 12, 6016 (2021).

5. Dettmers, C. et al. Comparison of regional cerebral blood flow with transcranial magnetic stimulation at different forces. J. Appl. Physiol. 81, 596–603 (1996).

6. Dai, T. H., Liu, J. Z., Sahgal, V., Brown, R. W. & Yue, G. H. Relationship between muscle output and functional MRI-measured brain activation. Exp. Brain Res. 140, 290–300 (2001).

7. Cramer, S. C. & Chopp, M. Recovery recapitulates ontogeny. Trends in Neurosciences 23, 265–271 (2000).

8. Joy, M. T. & Carmichael, S. T. Learning and stroke recovery: parallelism of biological substrates. Semin. Neurol. 41, 147–156 (2021).

9. Nudo, R. J. Recovery after brain injury: mechanisms and principles. Front. Hum. Neurosci. 7, 887 (2013).

10. Nakano, H., Tang, Y., Morita, T. & Naito, E. Theoretical proposal for restoration of hand motor function based on plasticity of motor-cortical interhemispheric interaction and its developmental rule. Front. Neurol. 15, 1408324 (2024).

11. Murphy, T. H. & Corbett, D. Plasticity during stroke recovery: from synapse to behaviour. Nat. Rev. Neurosci. 10, 861–872 (2009). (Review)

12. Li, W. et al. Changing genes, cells and networks to reprogram the brain after stroke. Nat. Neurosci. 28, 1130–1145 (2025). (Review)

13. Nato, G. et al. Brain injury reactivates a developmental program driving genesis and integration of transient LGE-class interneurons. Stem Cell Reports 21, 103015 (2026).

14. Magee, J. C. & Grienberger, C. Synaptic plasticity forms and functions. Annu. Rev. Neurosci. 43, 95–117 (2020).

15. Jang, H. J., McMahon Ward, R., Golden, C. E. M. & Constantinople, C. M. Acetylcholine demixes heterogeneous dopamine signals for learning and moving. Nature Neuroscience 29, 840–850 (2026).

16. Williams, B. M. et al. Neuromodulator timing regulates adult cortical plasticity via the synaptic eligibility trace. Brain Stimul. 19, 103004 (2026).

17. Gerstner, W., Lehmann, M., Liakoni, V., Corneil, D. & Brea, J. Eligibility traces and plasticity on behavioral time scales: experimental support of NeoHebbian three-factor learning rules. Front. Neural Circuits 12, 53–69 (2018).

18. Izhikevich, E. M. Solving the distal reward problem through linkage of STDP and dopamine signaling. BMC Neurosci. 8, S15 (2007).

19. Legenstein, R., Pecevski, D. & Maass, W. A learning theory for reward-modulated spike-timing-dependent plasticity with application to biofeedback. PLOS Comput. Biol. 4, e1000180 (2008).

20. Brzosko, Z., Schultz, S. & Paulsen, O. Retroactive modulation of spike timing-dependent plasticity by dopamine. eLife 4, e09685 (2015).

21. Bittner, K. C., Milstein, A. D., Grienberger, C., Romani, S. & Magee, J. C. Behavioral time scale synaptic plasticity underlies CA1 place fields. Science 357, 1033–1036 (2017).

22. Vendrell-Llopis, N. et al. Dopamine D1 receptor activation in the striatum is sufficient to drive reinforcement of anteceding cortical patterns. Neuron 113, 785–794.e9 (2025).

23. Yagishita, S. et al. A critical time window for dopamine actions on the structural plasticity of dendritic spines. Science 345, 1616–1620 (2014).

24. He, K. et al. Distinct eligibility traces for LTP and LTD in cortical synapses. Neuron 88, 528–538 (2015).

25. Hosp, J. A., Pekanovic, A., Rioult-Pedotti, M. S. & Luft, A. R. Dopaminergic projections from midbrain to primary motor cortex mediate motor skill learning. J. Neurosci. 31, 2481–2487 (2011).

26. Molina-Luna, K. et al. Dopamine in motor cortex is necessary for skill learning and synaptic plasticity. PLOS One 4, e7082 (2009).

27. Ghanayim, A. et al. VTA projections to M1 are essential for reorganization of layer 2-3 network dynamics underlying motor learning. Nat. Commun. 16, 200–217 (2025).

28. Leemburg, S., Canonica, T. & Luft, A. Motor skill learning and reward consumption differentially affect VTA activation. Sci. Rep. 8, 687–698 (2018).

29. Bova, A. et al. Precisely timed dopamine signals establish distinct kinematic representations of skilled movements. eLife 9, e61591 (2020).

30. Gorkowska-Nosal, M., Drwiega, G., Szumiec, L., Rodriguez Parkitna, J. & Cieslak, P. E. The role of dopamine-sensitive motor cortical circuits in the development and execution of skilled forelimb movements. iScience 29, 114983 (2026).

31. Redgrave, P. & Gurney, K. The short-latency dopamine signal: a role in discovering novel actions? Nat. Rev. Neurosci. 7, 967–975 (2006).

32. Schultz, W. Predictive reward signal of dopamine neurons. J. Neurophysiol. 80, 1–27 (1998).

33. Holzbaur, K. R. S., Delp, S. L., Gold, G. E. & Murray, W. M. Moment-generating capacity of upper limb muscles in healthy adults. J. Biomech. 40, 2442–2449 (2007).

34. Lieber, R. L., Jacobson, M. D., Fazeli, B. M., Abrams, R. A. & Botte, M. J. rchitecture of selected muscles of the arm and forearm: anatomy and implications for tendon transfer. J. Hand Surg. 17, 787–798 (1992).

35. Engelhard, B. et al. Specialized coding of sensory, motor and cognitive variables in VTA dopamine neurons. Nature 570, 509–513 (2019).

36. Greenstreet, F. et al. Dopaminergic action prediction errors serve as a value-free teaching signal. Nature 643, 1333–1342 (2025).

37. Glover, I. S. & Baker, S. N. Both corticospinal and reticulospinal tracts control force of contraction. J. Neurosci. 42, 3150–3164 (2022).

38. Danielson, T. L. et al. Activity in the pontine reticular nuclei scales with handgrip force in humans. J. Neurophysiol. 131, 807–814 (2024).

39. Akalu, Y. et al. Resistance training tempo selectively modulates corticospinal and reticulospinal excitability in humans. J. Physiol. 604, 3314–3343 (2026).

40. Basile, D. C., Paris, M. T., McNeil, C. J. & Rice, C. L. Investigating premotor corticospinal excitability in fast and slow voluntary contractions of the elbow flexors. J. Neurophysiol. 132, 1671–1679 (2024).

41. Roelfsema, P. R. & Holtmaat, A. Control of synaptic plasticity in deep cortical networks. Nat Rev Neurosci 19, 166–180 (2018).

42. Richards, B. A. & Lillicrap, T. P. Can neocortical feedback alter the sign of plasticity? Nat. Rev. Neurosci. 19, 636–636 (2018).

43. Guerguiev, J., Lillicrap, T. P. & Richards, B. A. Towards deep learning with segregated dendrites. eLife 6, e22901 (2017).

44. Williams, L. E. & Holtmaat, A. Higher-order thalamocortical inputs gate synaptic long-term potentiation via disinhibition. Neuron 101, 91–102.e4 (2019).

45. Dhawale, A. K., Smith, M. A. & Ölveczky, B. P. The Role of Variability in Motor Learning. Annu. Rev. Neurosci. 40, 479–498 (2017).

46. Ellis, M. D., Carmona, C., Drogos, J. & Dewald, J. P. A. Progressive abduction loading therapy with horizontal-plane viscous resistance targeting weakness and flexion synergy to treat upper limb function in chronic hemiparetic stroke: a randomized clinical trial. Front. Neurol. 9, 71–84 (2018).

47. Feydy, A. et al. Longitudinal study of motor recovery after stroke: recruitment and focusing of brain activation. Stroke 33, 1610–1617 (2002).

48. Li, S., Chen, Y.-T., Francisco, G. E., Zhou, P. & Rymer, W. Z. A unifying pathophysiological account for post-stroke spasticity and disordered motor control. Front. Neurol. 10, 468 (2019).

49. Zaaimi, B., Edgley, S. A., Soteropoulos, D. S. & Baker, S. N. Changes in descending motor pathway connectivity after corticospinal tract lesion in macaque monkey. Brain 135, 2277–2289 (2012).

50. Beer, R. F., Ellis, M. D., Holubar, B. G. & Dewald, J. P. A. Impact of gravity loading on post-stroke reaching and its relationship to weakness. Muscle Nerve 36, 242–250 (2007).

51. Sukal, T. M., Ellis, M. D. & Dewald, J. P. A. Shoulder abduction-induced reductions in reaching work area following hemiparetic stroke: neuroscientific implications. Exp. Brain Res. 183, 215–223 (2007).

52. Lan, Y., Yao, J. & Dewald, J. P. A. The Impact of Shoulder Abduction Loading on Volitional Hand Opening and Grasping in Chronic Hemiparetic Stroke. Neurorehabil Neural Repair 31, 521–529 (2017).

53. Rehme, A. K., Eickhoff, S. B., Wang, L. E., Fink, G. R. & Grefkes, C. Dynamic causal modeling of cortical activity from the acute to the chronic stage after stroke. Neuroimage 55, 1147–1158 (2011).

54. Rosso, C. et al. Contribution of corticospinal tract and functional connectivity in hand motor impairment after stroke. PLOS One 8, e73164 (2013).

55. Desowska, A. & Turner, D. L. Dynamics of brain connectivity after stroke. Rev. Neurosci. 30, 605–623 (2019).

56. Baldassarre, A. et al. Dynamic topological changes of the motor network after stroke. NeuroImage: Clin. 48, 103907 (2025).

57. Jiang, Z. et al. Dynamic reorganization of task-related network interactions in post-stroke aphasia recovery. Brain 148, 3563–3575 (2025).

58. Lüchinger, A. B., Hadders-Algra, M., Van Kan, C. M. & De Vries, J. I. P. Fetal onset of general movements. Pediatr. Res. 63, 191–195 (2008).

59. Hadders-Algra, M. General movements: a window for early identification of children at high risk for developmental disorders. J. Pediatr. 145, S12–S18 (2004).

60. De Vries, J. I. P., Visser, G. H. A. & Prechtl, H. F. R. The emergence of fetal behaviour. I. Qualitative aspects. Early Hum. Dev. 7, 301–322 (1982).

61. Trägenap, S., Whitney, D. E., Fitzpatrick, D. & Kaschube, M. The developmental emergence of reliable cortical representations. Nat. Neurosci. 28, 394–405 (2025).

62. McPherson, L. M. & Dewald, J. P. A. Abnormal synergies and associated reactions posthemiparetic stroke reflect muscle activation patterns of brainstem motor pathways. Front. Neurol. 13, 934670 (2022).

63. Grefkes, C. & Fink, G. R. Reorganization of cerebral networks after stroke: new insights from neuroimaging with connectivity approaches. Brain 134, 1264–1276 (2011).

64. Yu, P. et al. Neuroimaging of motor recovery after ischemic stroke − functional reorganization of motor network. NeuroImage: Clin. 43, 103636 (2024).

65. Sarkar, A. et al. Inter-individual variability in motor output is driven by recruitment gain in the corticospinal tract rather than motor threshold. Brain Sci. 12, 1401 (2022).

66. Scheidtmann, K., Fries, W., Müller, F. & Koenig, E. Effect of levodopa in combination with physiotherapy on functional motor recovery after stroke: a prospective, randomised, double-blind study. The Lancet 358, 787–790 (2001).

67. Acler, M., Fiaschi, A. & Manganotti, P. Long-term levodopa administration in chronic stroke patients. A clinical and neurophysiologic single-blind placebo-controlled cross-over pilot study. Restor. Neurol. Neurosci. 27, 277–283 (2009).

68. Sonde, L. & Lökk, J. Effects of amphetamine and/or l-dopa and physiotherapy after stroke? a blinded randomized study. Acta Neurol. Scand. 115, 55–59 (2007).

69. Ford, G. A. et al. Dopamine augmented rehabilitation in stroke (DARS): a multicentre double-blind, randomised controlled trial of co-careldopa compared with placebo, in addition to routine NHS occupational and physical therapy, delivered early after stroke on functional recovery. Effic. Mech. Eval. 6, 1–138 (2019).

70. Engelter, S. T. et al. Levodopa added to stroke rehabilitation: the ESTREL randomized clinical trial. JAMA 334, 1523–1532 (2025).

## References

1. Howe, M. W. & Dombeck, D. A. Rapid signalling in distinct dopaminergic axons during locomotion and reward. Nature 535, 505–510 (2016).

2. Vitrac, C. et al. Dopamine control of pyramidal neuron activity in the primary motor cortex via D2 receptors. Front. Neural Circuits 8, 13 (2014). doi:10.3389/fncir.2014.00013.

3. Groppa, S. et al. The human dorsal premotor cortex facilitates the excitability of ipsilateral primary motor cortex via a short latency cortico-cortical route. Hum. Brain Mapp. 33, 419–430 (2012).

4. Peters, A. J., Chen, S. X. & Komiyama, T. Emergence of reproducible spatiotemporal activity during motor learning. Nature 510, 263–267 (2014).

5. Masamizu, Y. et al. Two distinct layer-specific dynamics of cortical ensembles during learning of a motor task. Nat. Neurosci. 17, 987–994 (2014).

6. Zagha, E., Ge, X. & McCormick, D. A. Competing neural ensembles in motor cortex gate goal-directed motor output. Neuron 88, 565–577 (2015).

7. Jung, K. et al. An adaptive behavioral control motif mediated by cortical axo-axonic inhibition. Nat. Neurosci. 26, 1379–1393 (2023).

8. Li, Q. et al. Refinement of learned skilled movement representation in motor cortex deep output layer. Nat. Commun. 8, 15834 (2017).

9. Tallis, G. M. The moment generating function of the truncated multi-normal distribution. J. R. Stat. Soc. B 23(1), 223–229 (1961). doi:10.1111/j.2517-6161.1961.tb00408.x.

10. Sampford, M. R. Some Inequalities on Mill’s Ratio and Related Functions. The Annals of Mathematical Statistics 24, 130–132 (1953).

11. Mills, J. P. Table of the ratio: area to bounding ordinate, for any portion of normal curve. Biometrika 18, 395–400 (1927).

