## Supplementary material for "Selectivity Dilution in Three-Factor Learning: A Finite-Population Model with Potential Implications for Motor Development and Post-Stroke Motor Rehabilitation": code: Fig 1 BCD.py

```
"""
Case 1: Flexor-dominant network (positive total force)
Synaptic connection pool: 100 each of forces +9 (biceps), +6 (brachioradialis), -9 (triceps); 5000 of 0.
This case represents a network with net flexor bias; success probability increases
monotonically with the number of activated synapses (n).

Three-panel figure:
(A) Full view: marginal success probability vs. n
(B) Zoomed view: marginal success probability vs. n (n ≤ 200)
(C) Selectivity dilution: conditional flexor ratio (r/n) in successful trials vs. n (n ≤ 200)
    with P_success > 0.2 filtering, showing mean ± 95% CI across 20 independent runs.
"""

import numpy as np
import matplotlib.pyplot as plt
import matplotlib
from scipy import stats
import warnings
import time

warnings.filterwarnings('ignore')

# ------------------ Nature-style plotting settings ------------------
matplotlib.rcParams.update({
    'font.family': 'sans-serif',
    'font.sans-serif': ['Arial', 'Helvetica', 'AppleGothic'],
    'font.size': 8,
    'axes.labelsize': 9,
    'axes.titlesize': 10,
    'xtick.labelsize': 7,
    'ytick.labelsize': 7,
    'legend.fontsize': 7,
    'figure.titlesize': 11,
    'figure.dpi': 300,
    'savefig.dpi': 300,
    'savefig.bbox': 'tight',
    'axes.spines.top': False,
    'axes.spines.right': False,
    'axes.unicode_minus': False
})


def create_synapse_pool():
    """
    Create the pool of synaptic connections.
    Total synapses = 100 (biceps, +9) + 100 (brachioradialis, +6)
                    + 100 (triceps, -9) + 5000 (non-elbow, 0) = 5300.
    """
    forces = []
    forces.extend([9] * 100)    # biceps (strong flexor)
    forces.extend([6] * 100)    # brachioradialis (moderate flexor)
    forces.extend([-9] * 100)   # triceps (extensor)
    forces.extend([0] * 5000)   # non-elbow (task-irrelevant)
    return np.array(forces)


def normal_approximation(n, threshold, N=5300):
    """
    Normal approximation for success probability (finite population).
    N = total number of synapses = 5300.
    """
    pop_mean = (100*9 + 100*6 + 100*(-9)) / 5300
    var_9 = 100 * (9 - pop_mean)**2
    var_6 = 100 * (6 - pop_mean)**2
    var_neg9 = 100 * (-9 - pop_mean)**2
    var_0 = 5000 * (0 - pop_mean)**2
    pop_var = (var_9 + var_6 + var_neg9 + var_0) / 5300

    sample_mean = n * pop_mean
    if n == N:
        sample_var = 0
    else:
        fpc = (N - n) / (N - 1)
        sample_var = n * pop_var * fpc

    if sample_var <= 0:
        return 1.0 if sample_mean > threshold else 0.0

    z = (threshold + 0.5 - sample_mean) / np.sqrt(sample_var)
    prob = 1 - stats.norm.cdf(z)
    return max(0.0, min(1.0, prob))


# ------------------ Monte Carlo for selectivity dilution ------------------
def conditional_flexor_ratio_single_run(n, synapse_pool, threshold, num_trials=2000):
    """Target = flexors (+9 or +6). Returns mean flexor ratio (r/n) across successful trials."""
    N = len(synapse_pool)
    ratios = []
    for _ in range(num_trials):
        indices = np.random.choice(N, size=n, replace=False)
        activated = synapse_pool[indices]
        total_force = np.sum(activated)
        if total_force > threshold:
            flexor_count = np.sum((activated == 9) | (activated == 6))
            ratio = flexor_count / n
            ratios.append(ratio)
    if not ratios:
        return np.nan
    return np.mean(ratios)


def compute_conditional_curve_with_ci(n_vals, synapse_pool, threshold,
                                      n_runs=20, num_trials_per_run=2000):
    all_run_means = []
    for run_idx in range(n_runs):
        run_means = []
        for n in n_vals:
            if n > len(synapse_pool):
                n = len(synapse_pool)
            r = conditional_flexor_ratio_single_run(n, synapse_pool, threshold, num_trials_per_run)
            run_means.append(r)
        all_run_means.append(run_means)
        # print progress
        if (run_idx + 1) % 5 == 0:
            print(f"    Run {run_idx+1}/{n_runs} done for threshold {threshold}")
    all_run_means = np.array(all_run_means)
    mean_vals = np.nanmean(all_run_means, axis=0)
    sem_vals = np.nanstd(all_run_means, axis=0, ddof=1) / np.sqrt(n_runs)
    t_val = stats.t.ppf(0.975, n_runs - 1)
    ci_vals = sem_vals * t_val
    return mean_vals, ci_vals


# ------------------ Main plotting ------------------
def main():
    synapse_pool = create_synapse_pool()
    N_total = len(synapse_pool)
    K_flexors = 200
    global_baseline = K_flexors / N_total

    print(f"Total synapses: {N_total}, net total force: {np.sum(synapse_pool)}")
    print(f"Flexor proportion: {global_baseline:.4f}")

    thresholds = [2, 32]
    colors = ['#1f77b4', '#d62728']
    labels = ['R = 2', 'R = 32']

    # n ranges
    n_full = np.arange(1, N_total, 50)
    n_full = np.append(n_full, N_total)   # ensure last point is N_total
    n_zoom = np.arange(1, 201, 1)         # dense for zoom and dilution

    # ---------- 1. Monte Carlo for success probability (for scatter points) ----------
    key_n = sorted(set([1,2,5,10,20,30,50,100,200,300,500,
                        800,1000,1500,2000,3000,4000,5000,5300]))
    sim_results = {}
    print("Running Monte Carlo for success probability (num_trials=10000)...")
    for thr, lab in zip(thresholds, labels):
        print(f"  {lab}")
        probs = []
        for n in key_n:
            if n > N_total:
                n = N_total
            p = estimate_success_probability(n, synapse_pool, thr, num_trials=10000)
            probs.append(p)
        sim_results[thr] = (key_n.copy(), probs.copy())

    # Normal approximation for full curves
    approx_full = {thr: (n_full.copy(), [normal_approximation(n, thr) for n in n_full]) for thr in thresholds}
    approx_zoom = {thr: [normal_approximation(n, thr) for n in n_zoom] for thr in thresholds}

    # ---------- 2. Selectivity dilution (conditional flexor ratio) ----------
    print("\nRunning Monte Carlo for conditional flexor ratio (20 runs x 100 trials)...")
    dil_results = {}
    for thr in thresholds:
        print(f"  R={thr}")
        mean_vals, ci_vals = compute_conditional_curve_with_ci(n_zoom, synapse_pool, thr,
                                                                n_runs=20, num_trials_per_run=100)
        dil_results[thr] = (mean_vals, ci_vals)

    # Precompute P_success for filtering dilution (using normal approximation)
    prob_success_2 = [normal_approximation(n, 2) for n in n_zoom]
    prob_success_32 = [normal_approximation(n, 32) for n in n_zoom]

    # ---------- 3. Plotting ----------
    fig, axes = plt.subplots(1, 3, figsize=(10.0, 3.5))  # 3 columns

    # ---- Panel 1: Full view ----
    ax1 = axes[0]
    for thr, col, lab in zip(thresholds, colors, labels):
        n_vals, probs = approx_full[thr]
        ax1.plot(n_vals, probs, color=col, lw=1.5, alpha=0.8, label=lab)
        n_sim, p_sim = sim_results[thr]
        ax1.scatter(n_sim, p_sim, color=col, s=20, alpha=0.8, edgecolors='k', linewidth=0.5, zorder=3)
    ax1.set_xlabel('Number of co-activated neurons (n)')
    ax1.set_ylabel('Probability of success, $P_{\\mathrm{success}}$')
    ax1.set_title('(B) Full view', fontweight='bold')
    ax1.set_xlim(0, 5400)
    ax1.set_ylim(-0.05, 1.05)
    ax1.legend(loc='lower right', frameon=False)

    # ---- Panel 2: Zoomed view (n≤200) ----
    ax2 = axes[1]
    for thr, col, lab in zip(thresholds, colors, labels):
        probs = approx_zoom[thr]
        ax2.plot(n_zoom, probs, color=col, lw=1.5, alpha=0.8, label=lab)
        n_sim, p_sim = sim_results[thr]
        zoom_mask = [n <= 200 for n in n_sim]
        ax2.scatter([n for n, m in zip(n_sim, zoom_mask) if m],
                    [p for p, m in zip(p_sim, zoom_mask) if m],
                    color=col, s=20, alpha=0.8, edgecolors='k', linewidth=0.5)
    ax2.set_xlabel('Number of co-activated neurons (n)')
    ax2.set_ylabel('$P_{\\mathrm{success}}$')
    ax2.set_title('(C) Zoomed view (n ≤ 200)', fontweight='bold')
    ax2.set_xlim(0, 205)
    ax2.set_ylim(-0.05, 1.05)
    ax2.legend(loc='lower right', frameon=False)

    # ---- Panel 3: Selectivity dilution ----
    ax3 = axes[2]
    # Compute marginal P_success for filtering (using normal approximation)
    prob_success_dict = {2: prob_success_2, 32: prob_success_32}
    for thr, col, lab in zip(thresholds, colors, labels):
        mean_vals, ci_vals = dil_results[thr]
        prob_success = prob_success_dict[thr]
        # Mask: not NaN and P_success > 0.2
        mask = (~np.isnan(mean_vals)) & (np.array(prob_success) > 0.2)
        n_vals = n_zoom[mask]
        means = mean_vals[mask]
        cis = ci_vals[mask]
        if len(n_vals) > 0:
            ax3.plot(n_vals, means, color=col, lw=1.5, alpha=0.8, label=lab)
            ax3.fill_between(n_vals, means - cis, means + cis,
                             color=col, alpha=0.2, edgecolor='none')

    # Global baseline
    ax3.axhline(y=global_baseline, color='gray', linestyle='--', lw=1.0,
                alpha=0.7, label=f'Baseline ({global_baseline:.2f})')
    # Mark P_success=0.2 boundary for R=2 (if exists)
    # Find first n where P_success drops below 0.2 for R=2
    for thr in [2]:
        if thr == 2:
            probs = [normal_approximation(n, thr) for n in n_zoom]
            boundary = None
            for idx, p in enumerate(probs):
                if p <= 0.2:
                    boundary = n_zoom[idx]
                    break
            if boundary is not None:
                ax3.axvline(x=boundary, color='gray', linestyle=':', lw=1.0, alpha=0.6,
                            label=f'P_success=0.2 boundary')
    ax3.set_xlabel('Number of co-activated neurons (n)')
    ax3.set_ylabel('Flexor ratio in successful trials (r / n)')
    ax3.set_title('(D) Selectivity dilution (n ≤ 200)', fontweight='bold')
    ax3.set_xlim(0, 205)
    ax3.set_ylim(-0.05, 1.05)
    ax3.legend(loc='upper right', frameon=False)

    # Adjust layout
    plt.tight_layout()
    plt.savefig('Fig1BCD_Flexor_Dominant.pdf', dpi=300, bbox_inches='tight')
    plt.savefig('Fig1BCD_Flexor_Dominant.svg', bbox_inches='tight')
    print("\nFigures saved as 'Fig1BCD_Flexor_Dominant.pdf' and .svg")
    plt.show()


# Helper: Monte Carlo for success probability (copied from original)
def simulate_trial(n, synapse_pool, threshold):
    indices = np.random.choice(len(synapse_pool), size=n, replace=False)
    total_force = np.sum(synapse_pool[indices])
    return total_force > threshold

def estimate_success_probability(n, synapse_pool, threshold, num_trials=100):
    successes = 0
    for _ in range(num_trials):
        if simulate_trial(n, synapse_pool, threshold):
            successes += 1
    return successes / num_trials


if __name__ == "__main__":
    np.random.seed(32)
    main()
```
