## Supplementary material for "Selectivity Dilution in Three-Factor Learning: A Finite-Population Model with Potential Implications for Motor Development and Post-Stroke Motor Rehabilitation": code: Fig 1 EFG.py

```
"""
Case: Extensor target in a flexor-dominant network (net negative total force)
Synaptic connection pool: 100 each of +9 (extensor), -6 (moderate flexor), -9 (strong flexor); 5000 of 0.
Success probability peaks at intermediate n and declines at high n.

Three-panel figure:
(A) Full view: marginal success probability vs. n
(B) Zoomed view: marginal success probability vs. n (n ≤ 200)
(C) Selectivity dilution: conditional proportion of +9 extensor neurons in successful trials vs. n (n ≤ 200)
    with P_success > 0.2 filtering, mean ± 95% CI across 20 runs, 100 trials per run per n.


    Note on the Discrepancy between Monte Carlo Scatters and Normal Approximation:
    In the low-n regime (n < 30, corresponding to the Left Edge), the Monte Carlo
    scatters appear systematically below the normal approximation curve.
    This is NOT a coding error or approximation failure, but a mathematical
    consequence of the extreme skewness of the finite-population hypergeometric
    distribution at small sample sizes.
    At the Left Edge, successful trials are driven not by mean-field force (which
    is insufficient to overcome R), but by rare, high-impact variance-driven
    fluctuations (sampling a few large +9 or +6 synapses). The normal approximation
    (CLT) over-smooths this tail probability, while the exact Monte Carlo estimates
    accurately capture the true success probability. Intriguingly, this inherent
    reliance on "variance-driven success" is precisely the mechanistic basis of
    the Left Edge strategy, as it confines reinforcement to a highly selective
    synaptic subset, thus circumventing the dilution constraint.

"""

import numpy as np
import matplotlib.pyplot as plt
import matplotlib
from scipy import stats
import warnings
import time

warnings.filterwarnings('ignore')

matplotlib.rcParams.update({
    'font.family': 'sans-serif',
    'font.sans-serif': ['Arial', 'Helvetica', 'AppleGothic'],
    'font.size': 8,
    'axes.labelsize': 9,
    'axes.titlesize': 10,
    'xtick.labelsize': 7,
    'ytick.labelsize': 7,
    'legend.fontsize': 7,
    'figure.titlesize': 11,
    'figure.dpi': 300,
    'savefig.dpi': 300,
    'savefig.bbox': 'tight',
    'axes.spines.top': False,
    'axes.spines.right': False,
    'axes.unicode_minus': False
})


def create_synapse_pool():
    """
    Pool: 100 each of +9 (extensor target), -6 (moderate flexor), -9 (strong flexor); 5000 zero.
    Total N = 5300.
    """
    forces = []
    forces.extend([9] * 100)    # extensor (target)
    forces.extend([-6] * 100)   # moderate flexor
    forces.extend([-9] * 100)   # strong flexor
    forces.extend([0] * 5000)   # irrelevant
    return np.array(forces)


def simulate_trial(n, synapse_pool, threshold):
    indices = np.random.choice(len(synapse_pool), size=n, replace=False)
    total_force = np.sum(synapse_pool[indices])
    return total_force > threshold


def estimate_probability(n, synapse_pool, threshold, num_trials=10000):
    successes = 0
    for _ in range(num_trials):
        if simulate_trial(n, synapse_pool, threshold):
            successes += 1
    return successes / num_trials


def normal_approximation_extensor(n, threshold, N=5300):
    """Normal approximation for marginal P_success in extensor-target case."""
    pop_mean = (100*9 + 100*(-6) + 100*(-9)) / N
    var_9 = 100 * (9 - pop_mean)**2
    var_neg6 = 100 * (-6 - pop_mean)**2
    var_neg9 = 100 * (-9 - pop_mean)**2
    var_0 = 5000 * (0 - pop_mean)**2
    pop_var = (var_9 + var_neg6 + var_neg9 + var_0) / N

    sample_mean = n * pop_mean
    if n == N:
        sample_var = 0
    else:
        fpc = (N - n) / (N - 1)
        sample_var = n * pop_var * fpc

    if sample_var <= 0:
        return 1.0 if sample_mean > threshold else 0.0

    z = (threshold + 0.5 - sample_mean) / np.sqrt(sample_var)
    prob = 1 - stats.norm.cdf(z)
    return max(0.0, min(1.0, prob))


# ------------------ 选择性稀释相关函数 ------------------
def conditional_target_ratio_single_run(n, synapse_pool, threshold, num_trials=100):
    """Target = +9 extensor neurons. Returns mean target ratio (r/n) across successful trials."""
    N = len(synapse_pool)
    ratios = []
    for _ in range(num_trials):
        indices = np.random.choice(N, size=n, replace=False)
        activated = synapse_pool[indices]
        total_force = np.sum(activated)
        if total_force > threshold:
            target_count = np.sum(activated == 9)
            ratio = target_count / n
            ratios.append(ratio)
    if not ratios:
        return np.nan
    return np.mean(ratios)


def compute_conditional_curve_with_ci(n_vals, synapse_pool, threshold,
                                      n_runs=20, num_trials_per_run=100):
    all_run_means = []
    for run_idx in range(n_runs):
        run_means = []
        for n in n_vals:
            if n > len(synapse_pool):
                n = len(synapse_pool)
            r = conditional_target_ratio_single_run(n, synapse_pool, threshold, num_trials_per_run)
            run_means.append(r)
        all_run_means.append(run_means)
        print(f"    Run {run_idx+1}/{n_runs} done for threshold {threshold}")
    all_run_means = np.array(all_run_means)
    mean_vals = np.nanmean(all_run_means, axis=0)
    sem_vals = np.nanstd(all_run_means, axis=0, ddof=1) / np.sqrt(n_runs)
    t_val = stats.t.ppf(0.975, n_runs - 1)
    ci_vals = sem_vals * t_val
    return mean_vals, ci_vals


def main():
    pool = create_synapse_pool()
    N_total = len(pool)
    K_target = 100  # +9 extensor neurons
    global_baseline = K_target / N_total

    print(f"Total synapses: {N_total}, net total force: {np.sum(pool)}")
    print(f"Target (+9 extensor) proportion: {global_baseline:.4f}")

    thresholds = [2, 32]
    colors = ['#1f77b4', '#d62728']
    labels = ['R = 2', 'R = 32']

    # n ranges
    n_full = np.arange(1, N_total, 50)
    n_full = np.append(n_full, N_total)   # ensure last point is N_total
    n_dense = np.arange(1, 201, 2)        # for zoom success probability
    n_dilution = np.arange(1, 201, 1)     # dense for dilution

    # ---------- 1. 蒙特卡洛模拟成功概率（散点） ----------
    key_n = sorted(set([1,2,5,10,20,30,50,100,200,300,500,
                        800,1000,1500,2000,3000,4000,5000,5300]))
    print("Running Monte Carlo for success probability (num_trials=10000)...")
    sim_results = {}
    for thr, lab in zip(thresholds, labels):
        print(f"  {lab}")
        probs = []
        for n in key_n:
            if n > N_total:
                n = N_total
            p = estimate_probability(n, pool, thr, num_trials=10000)
            probs.append(p)
        sim_results[thr] = (key_n.copy(), probs.copy())

    # 正态近似曲线
    approx_full = {thr: (n_full.copy(), [normal_approximation_extensor(n, thr, N_total) for n in n_full]) for thr in thresholds}
    approx_dense = {thr: [normal_approximation_extensor(n, thr, N_total) for n in n_dense] for thr in thresholds}

    # ---------- 2. 选择性稀释（20 runs × 100 trials per n） ----------
    print("\nRunning Monte Carlo for conditional target ratio (20 runs × 100 trials)...")
    dil_results = {}
    for thr in thresholds:
        print(f"  R={thr}")
        mean_vals, ci_vals = compute_conditional_curve_with_ci(
            n_dilution, pool, thr, n_runs=20, num_trials_per_run=100
        )
        dil_results[thr] = (mean_vals, ci_vals)

    # 预计算 P_success 用于过滤稀释
    prob_success_2 = [normal_approximation_extensor(n, 2, N_total) for n in n_dilution]
    prob_success_32 = [normal_approximation_extensor(n, 32, N_total) for n in n_dilution]
    prob_success_dict = {2: prob_success_2, 32: prob_success_32}

    # ---------- 3. 绘图 ----------
    fig, axes = plt.subplots(1, 3, figsize=(10.0, 3.5))  # 三列

    # ---- Panel A: 全览 ----
    ax1 = axes[0]
    for thr, col, lab in zip(thresholds, colors, labels):
        n_vals, probs = approx_full[thr]
        ax1.plot(n_vals, probs, color=col, lw=1.5, alpha=0.8, label=lab)
        n_sim, p_sim = sim_results[thr]
        ax1.scatter(n_sim, p_sim, color=col, s=20, alpha=0.8, edgecolors='k', linewidth=0.5, zorder=3)
    ax1.set_xlabel('Number of co-activated neurons (n)')
    ax1.set_ylabel('Probability of success, $P_{\\mathrm{success}}$')
    ax1.set_title('(E) Full view', fontweight='bold')
    ax1.set_xlim(0, 5400)
    ax1.set_ylim(-0.05, 1.05)
    ax1.legend(loc='upper right', frameon=False)

    # ---- Panel B: 放大 (n ≤ 200) ----
    ax2 = axes[1]
    for thr, col, lab in zip(thresholds, colors, labels):
        probs = approx_dense[thr]
        ax2.plot(n_dense, probs, color=col, lw=1.5, alpha=0.8, label=lab)
        n_sim, p_sim = sim_results[thr]
        zoom_mask = [n <= 200 for n in n_sim]
        ax2.scatter([n for n, m in zip(n_sim, zoom_mask) if m],
                    [p for p, m in zip(p_sim, zoom_mask) if m],
                    color=col, s=20, alpha=0.8, edgecolors='k', linewidth=0.5)
    ax2.set_xlabel('Number of co-activated neurons (n)')
    ax2.set_ylabel('$P_{\\mathrm{success}}$')
    ax2.set_title('(F) Zoomed view (n ≤ 200)', fontweight='bold')
    ax2.set_xlim(0, 201)
    ax2.set_ylim(-0.05, 1.05)
    ax2.legend(loc='upper right', frameon=False)

    # ---- Panel C: 选择性稀释 ----
    ax3 = axes[2]
    for thr, col, lab in zip(thresholds, colors, labels):
        mean_vals, ci_vals = dil_results[thr]
        prob_success = prob_success_dict[thr]
        # 过滤: P_success > 0.2 且 非 NaN
        mask = (~np.isnan(mean_vals)) & (np.array(prob_success) > 0.2)
        n_vals = n_dilution[mask]
        means = mean_vals[mask]
        cis = ci_vals[mask]
        if len(n_vals) > 0:
            ax3.plot(n_vals, means, color=col, lw=1.5, alpha=0.8, label=lab)
            ax3.fill_between(n_vals, means - cis, means + cis,
                             color=col, alpha=0.2, edgecolor='none')

    ax3.axhline(y=global_baseline, color='gray', linestyle='--', lw=1.0,
                alpha=0.7, label=f'Baseline ({global_baseline:.2f})')
    # 标记 R=2 的 P_success=0.2 边界（左右边界）
    for thr in [2]:
        probs = prob_success_dict[thr]
        left_idx = None
        right_idx = None
        for i, p in enumerate(probs):
            if p > 0.2:
                left_idx = i
                break
        for i in range(len(probs)-1, -1, -1):
            if probs[i] > 0.2:
                right_idx = i
                break
        if left_idx is not None and right_idx is not None:
            ax3.axvline(x=n_dilution[left_idx], color='gray', linestyle=':', lw=1.0, alpha=0.6,
                        label='P_success=0.2 boundary')
            ax3.axvline(x=n_dilution[right_idx], color='gray', linestyle=':', lw=1.0, alpha=0.6)
    ax3.set_xlabel('Number of co-activated neurons (n)')
    ax3.set_ylabel('S(n) (extensor training)')
    ax3.set_title('(G) Selectivity dilution (n ≤ 200)', fontweight='bold')
    ax3.set_xlim(0, 201)
    ax3.set_ylim(-0.05, 1.05)
    ax3.legend(loc='upper right', frameon=False)

    plt.tight_layout()
    plt.savefig('Fig1EFG_Extensor_Target.pdf', dpi=300, bbox_inches='tight')
    plt.savefig('Fig1EFG_Extensor_Target.svg', bbox_inches='tight')
    print("\nFigures saved as 'Fig1EFG_Extensor_Target.pdf' and .svg")
    plt.show()


if __name__ == "__main__":
    np.random.seed(32)
    main()
```
