## Supplementary material for "Selectivity Dilution in Three-Factor Learning: A Finite-Population Model with Potential Implications for Motor Development and Post-Stroke Motor Rehabilitation": code: Fig 2 A-D.py

```
"""
Extensor strengthening in flexor-dominant pool (R=2 vs R=12)
Trends in Neurosciences style with smoothing
- Static flexor bias via asymmetric force distribution
- brachioradialis force changed from -2 to -6 (moderate flexor)
- GMA-modulated sampling + failure-dependent depression
- Demonstrates narrow window for selective extensor reinforcement
- Plot: 2x2 subplots (composition + force/GMA for each R)

Ensemble-Based Reinforcement Learning with Structural Plasticity

This implementation models motor skill learning using a discrete, ensemble-based
selection mechanism rather than continuous weight updates. The algorithm operates
on three core principles:

1. Stochastic Recruitment: At each time step, a subset of synapses (the activated
   ensemble) is sampled probabilistically from the total pool, with bias toward
   previously reinforced connections.

2. Success-Gated Reinforcement (Three-Factor Rule): If the generated torque exceeds
   the resistance threshold (F > R), the activated ensemble is added to the
   functional pool (reinforced set). This mirrors the dopaminergic third factor in
   biological three-factor learning: success gates plasticity.

3. Competitive Pruning: The functional pool is maintained at a fixed size via
   random removal of excess synapses—a form of homeostatic competition that
   preserves efficiency while preventing runaway potentiation.

This discrete ensemble approach confers several advantages over conventional
continuous-weight models: (i) computational sparsity, as updates operate only on
the activated subset; (ii) inherent robustness to catastrophic interference, as
knowledge is stored in binary membership rather than scalar weights; and
(iii) alignment with structural plasticity observed in motor learning, where skill
acquisition correlates with the formation and refinement of task-specific synaptic
ensembles.

The simulation parameters (resistance R, GMA dynamics, pool sizes) are chosen to
illustrate the theoretical framework described in the accompanying manuscript,
particularly the concept of "selectivity dilution" under high modulatory drive.
"""

import random
import numpy as np
from collections import Counter
import matplotlib.pyplot as plt
from scipy import stats

# ------------------ Muscle mapping (brachioradialis force = -6) ------------------
FORCE_TO_MUSCLE = {
    9:   'triceps',          # extensor
    -9:  'biceps',           # strong flexor
    -6:  'brachioradialis',  # moderate flexor (was -6)
    0:   'non-elbow'
}
MUSCLE_ORDER = ['biceps', 'brachioradialis', 'triceps', 'non-elbow']
MUSCLE_LABELS = {
    'biceps': 'Biceps',
    'brachioradialis': 'Brachioradialis',
    'triceps': 'Triceps',
    'non-elbow': 'Non-elbow'
}
# Color-blind friendly palette (Okabe-Ito)
MUSCLE_COLORS = {
    'biceps': '#D55E00',          # orange
    'brachioradialis': '#0072B2', # blue
    'triceps': '#CC79A7',         # pinkish purple
    'non-elbow': '#999999'        # grey
}

# ------------------ Circuit class ------------------
class ExtensorCircuitWithDepression:
    def __init__(self, R=12.0, base_decay=0.1, min_decay=0.01, max_gma=20):
        self.R = R
        self.base_decay = base_decay
        self.min_decay = min_decay
        self.max_gma = max_gma
        self.target_size = 100

        # Neuron pools: brachioradialis force changed to -6
        self.neuron_props = {
            'flex_strong':  {'count': 100, 'force': -9},
            'flex_mod':     {'count': 100, 'force': -6},   # was -2
            'non_elbow':    {'count': 5000, 'force': 0},
            'extensor':     {'count': 100, 'force': 9}
        }
        self._init_pools()
        self.history = {'force': [], 'gma': [], 'extensor_ratio': [],
                        'reinforced_composition': []}

    def _init_pools(self):
        self.all_neurons = []
        for p in self.neuron_props.values():
            self.all_neurons.extend([p['force']] * p['count'])
        self.reinforced = random.sample(self.all_neurons, self.target_size)
        self.non_reinforced = self._subtract(self.all_neurons, self.reinforced)

    def _subtract(self, a, b):
        ca, cb = Counter(a), Counter(b)
        for k, v in cb.items():
            if ca[k] < v:
                raise ValueError('subset error')
        return list((ca - cb).elements())

    def _safe_remove(self, lst, to_remove):
        cnt = Counter(to_remove)
        new = []
        for x in lst:
            if cnt[x] > 0:
                cnt[x] -= 1
            else:
                new.append(x)
        return new

    def _activation(self, gma):
        spec_size = 15 + int(gma * 0.01 * len(self.reinforced))
        spec_size = min(spec_size, len(self.reinforced))
        spec = random.sample(self.reinforced, spec_size) if spec_size > 0 else []
        if gma > 0:
            nonspec_size = int(gma * 0.001 * len(self.non_reinforced))
            if nonspec_size > 0 and len(self.non_reinforced) > 0:
                nonspec_size = min(nonspec_size, len(self.non_reinforced))
                nonspec = random.sample(self.non_reinforced, nonspec_size)
            else:
                nonspec = []
        else:
            nonspec = []
        return spec + nonspec

    def _reinforce(self, activated):
        self.reinforced.extend(activated)
        self.non_reinforced = self._safe_remove(self.non_reinforced, activated)

    def _depress(self, activated, gma):
        rate = self.min_decay + (gma / self.max_gma) * (self.base_decay - self.min_decay)
        active_in_reinf = [x for x in activated if x in self.reinforced]
        n_remove = int(len(active_in_reinf) * rate)
        if n_remove > 0 and active_in_reinf:
            max_removable = max(0, len(self.reinforced) - 1)
            n_remove = min(n_remove, max_removable, len(active_in_reinf))
            if n_remove > 0:
                to_remove = random.sample(active_in_reinf, n_remove)
                self.reinforced = self._safe_remove(self.reinforced, to_remove)
                self.non_reinforced.extend(to_remove)

    def _prune(self):
        if len(self.reinforced) > self.target_size:
            excess = len(self.reinforced) - self.target_size
            indices_to_remove = random.sample(range(len(self.reinforced)), excess)
            for idx in sorted(indices_to_remove, reverse=True):
                pruned = self.reinforced.pop(idx)
                self.non_reinforced.append(pruned)

    def simulate(self, max_trials=500):
        gma = 0
        consec = 0
        stab_step = None
        for step in range(max_trials):
            activated = self._activation(gma)
            F = sum(activated)

            total = len(self.reinforced)
            if total == 0:
                step_composition = {muscle: 0.0 for muscle in FORCE_TO_MUSCLE.values()}
            else:
                comp = Counter(self.reinforced)
                step_composition = {muscle: comp.get(force, 0) / total
                                     for force, muscle in FORCE_TO_MUSCLE.items()}
            self.history['reinforced_composition'].append(step_composition)

            if F > self.R:
                self._reinforce(activated)
                self._prune()
                consec += 1
                gma = max(0, gma - 2)
                if consec >= 10 and stab_step is None:
                    stab_step = step
            else:
                self._depress(activated, gma)
                consec = 0
                gma = min(gma + 1, self.max_gma)

            self.history['force'].append(F)
            self.history['gma'].append(gma)
            ext_count = sum(1 for x in self.reinforced if x == 9)
            ext_ratio = ext_count / len(self.reinforced) if self.reinforced else 0
            self.history['extensor_ratio'].append(ext_ratio)

        return stab_step, self.history

# ------------------ Batch simulation ------------------
def run_experiment(R, n_runs=20, max_trials=500):
    all_compositions = []
    all_force_histories = []
    all_gma_histories = []
    stable_count = 0
    stab_trials = []

    for run in range(n_runs):
        model = ExtensorCircuitWithDepression(R=R, base_decay=0.1, min_decay=0.01)
        sf, hist = model.simulate(max_trials)
        all_compositions.append(hist['reinforced_composition'])
        all_force_histories.append(hist['force'])
        all_gma_histories.append(hist['gma'])
        if sf is not None:
            stable_count += 1
            stab_trials.append(sf)

    if stab_trials:
        avg_step = np.mean(stab_trials)
        print(f"R={R}: {stable_count}/{n_runs} stabilized, avg step {avg_step:.1f}")
    else:
        print(f"R={R}: {stable_count}/{n_runs} stabilized")

    # Compute statistics for muscle composition
    n_muscles = len(MUSCLE_ORDER)
    data = np.zeros((n_runs, max_trials, n_muscles))
    for r, comp_list in enumerate(all_compositions):
        for t, comp_dict in enumerate(comp_list):
            for m_idx, muscle in enumerate(MUSCLE_ORDER):
                data[r, t, m_idx] = comp_dict.get(muscle, 0.0)

    mean = np.mean(data, axis=0)
    sem = stats.sem(data, axis=0)
    ci = sem * stats.t.ppf(0.975, n_runs - 1)

    trials = list(range(max_trials))
    stats_dict = {}
    for m_idx, muscle in enumerate(MUSCLE_ORDER):
        stats_dict[muscle] = {
            'mean': mean[:, m_idx],
            'ci': ci[:, m_idx]
        }

    return trials, stats_dict, all_compositions, all_force_histories, all_gma_histories

# ------------------ Unified plotting with smoothing ------------------
def plot_combined(stats_R2, stats_R12, output_base='Extensor_Strengthening_Combined'):

    # 平滑函数：滑动窗口平均，窗口大小 9
    def smooth_trace(data, window=9):
        if len(data) < window:
            return data
        kernel = np.ones(window) / window
        smoothed = np.convolve(data, kernel, mode='same')
        # 边缘修正，防止两端被下拉
        pad = window // 2
        smoothed[:pad] = data[:pad]
        smoothed[-pad:] = data[-pad:]
        return smoothed

    plt.rcParams.update({
        'font.family': 'sans-serif',
        'font.sans-serif': ['Arial', 'Helvetica'],
        'font.size': 7,
        'axes.labelsize': 8,
        'axes.titlesize': 9,
        'axes.spines.top': False,
        'axes.spines.right': False,
        'axes.linewidth': 0.8,
        'xtick.direction': 'in',
        'ytick.direction': 'in',
        'legend.fontsize': 6,
        'figure.dpi': 300,
        'savefig.dpi': 300,
        'savefig.bbox': 'tight'
    })

    fig, axes = plt.subplots(2, 2, figsize=(7.2, 5.5))
    R_values = [2, 12]
    all_stats = [stats_R2, stats_R12]
    all_compositions = [stats_R2[2], stats_R12[2]]
    all_force = [stats_R2[3], stats_R12[3]]
    all_gma = [stats_R2[4], stats_R12[4]]

    for row, (R, stats_data) in enumerate(zip(R_values, all_stats)):
        trials, stats_dict, comps, forces, gmas = stats_data

        # -------- Left subplot: composition --------
        ax_comp = axes[row, 0]
        # Plot first 5 single runs (translucent, NO smoothing)
        for run_idx in range(min(5, len(comps))):
            run_data = comps[run_idx]
            for muscle in MUSCLE_ORDER:
                trace = [d.get(muscle, 0.0) for d in run_data]
                ax_comp.plot(trials, trace, color=MUSCLE_COLORS[muscle],
                             lw=0.2, alpha=0.15, rasterized=True)
        # Plot mean ± CI (mean smoothed)
        for muscle in MUSCLE_ORDER:
            mean_raw = stats_dict[muscle]['mean']
            ci_raw = stats_dict[muscle]['ci']
            mean_smooth = smooth_trace(mean_raw, window=9)
            ax_comp.plot(trials, mean_smooth, color=MUSCLE_COLORS[muscle],
                         lw=1.2, label=MUSCLE_LABELS[muscle])
            ax_comp.fill_between(trials, mean_raw - ci_raw, mean_raw + ci_raw,
                                 color=MUSCLE_COLORS[muscle], alpha=0.2)

        ax_comp.set_xlabel('Time (trials)')
        ax_comp.set_ylabel('Proportion in\nreinforced pool')
        ax_comp.set_title(f'R = {R}', fontweight='bold')
        ax_comp.set_ylim(0, 1)
        ax_comp.spines['top'].set_visible(False)
        ax_comp.spines['right'].set_visible(False)

        # -------- Right subplot: Force and GMA (twin y) --------
        ax_force = axes[row, 1]
        # Compute mean ± CI for force
        force_array = np.array([h[:len(trials)] for h in forces])
        f_mean = np.mean(force_array, axis=0)
        f_sem = stats.sem(force_array, axis=0)
        f_ci = f_sem * stats.t.ppf(0.975, force_array.shape[0] - 1)
        # GMA
        gma_array = np.array([h[:len(trials)] for h in gmas])
        g_mean = np.mean(gma_array, axis=0)
        g_sem = stats.sem(gma_array, axis=0)
        g_ci = g_sem * stats.t.ppf(0.975, gma_array.shape[0] - 1)

        # Smooth the mean traces
        f_mean_smooth = smooth_trace(f_mean, window=9)
        g_mean_smooth = smooth_trace(g_mean, window=9)

        color_force = '#1f77b4'
        color_gma = '#ff7f0e'
        # Force (left y)
        ax_force.fill_between(trials, f_mean - f_ci, f_mean + f_ci,
                              color=color_force, alpha=0.2)
        ax_force.plot(trials, f_mean_smooth, color=color_force, lw=1.2, label='Force')
        ax_force.axhline(y=R, color='red', linestyle='--', lw=0.8, label='Threshold')
        ax_force.set_xlabel('Time (trials)')
        ax_force.set_ylabel('Summed force (a.u.)', color=color_force)
        ax_force.tick_params(axis='y', labelcolor=color_force)
        ax_force.spines['top'].set_visible(False)
        ax_force.spines['right'].set_visible(False)

        # GMA (right y)
        ax_gma = ax_force.twinx()
        ax_gma.fill_between(trials, g_mean - g_ci, g_mean + g_ci,
                            color=color_gma, alpha=0.2)
        ax_gma.plot(trials, g_mean_smooth, color=color_gma, lw=1.2, label='GMA')
        ax_gma.set_ylabel('GMA', color=color_gma)
        ax_gma.tick_params(axis='y', labelcolor=color_gma)
        ax_gma.spines['top'].set_visible(False)
        ax_gma.spines['right'].set_visible(False)

        # Combine legends
        lines1, labels1 = ax_force.get_legend_handles_labels()
        lines2, labels2 = ax_gma.get_legend_handles_labels()
        ax_force.legend(lines1 + lines2, labels1 + labels2,
                        loc='upper right', frameon=False, fontsize=6)

    # Overall figure legend for muscles (place outside or at top)
    handles = [plt.Line2D([0], [0], color=MUSCLE_COLORS[m], lw=2) for m in MUSCLE_ORDER]
    fig.legend(handles, [MUSCLE_LABELS[m] for m in MUSCLE_ORDER],
               loc='upper center', bbox_to_anchor=(0.5, 1.02),
               ncol=4, frameon=False, fontsize=7)

    plt.tight_layout()
    plt.subplots_adjust(top=0.92)
    for ext in ['pdf', 'svg']:
        plt.savefig(f"{output_base}.{ext}", dpi=300, bbox_inches='tight')
    plt.close()
    print(f"Saved {output_base}.pdf and .svg")

# ------------------ Main ------------------
if __name__ == "__main__":
    random.seed(32)
    np.random.seed(32)

    print("Running R=2 ...")
    stats_R2 = run_experiment(R=2, n_runs=20, max_trials=500)
    print("Running R=12 ...")
    stats_R12 = run_experiment(R=12, n_runs=20, max_trials=500)

    plot_combined(stats_R2, stats_R12, output_base='A Extensor_Strengthening_Combined')
```
