## Supplementary material for "Selectivity Dilution in Three-Factor Learning: A Finite-Population Model with Potential Implications for Motor Development and Post-Stroke Motor Rehabilitation": code: Fig 2 G-J Σ╣ïσÉÄσê╗μäÅσó₧σèáGMA.py

```
"""
Modified version of Extensor_Strengthening_R2_R12 to test the effect of forced high GMA
after an extensor-dominant circuit has already been established (step >= 400).
- Original R=2 (no GMA floor) vs Modified R=2 (GMA lower bound = 10 after step 400)
- Demonstrates that artificially increasing GMA can degrade extensor performance.
- Plot: 2x2 subplots with GMA color #ff7f0e (orange) matching original Fig2A.py style.
- Labels: 'Constraint onset' and 'GMA lower bound' for academic clarity.
"""

import random
import numpy as np
from collections import Counter
import matplotlib.pyplot as plt
from scipy import stats

# ------------------ Muscle mapping (brachioradialis force = -6) ------------------
FORCE_TO_MUSCLE = {
    9:   'triceps',          # extensor
    -9:  'biceps',           # strong flexor
    -6:  'brachioradialis',  # moderate flexor
    0:   'non-elbow'
}
MUSCLE_ORDER = ['biceps', 'brachioradialis', 'triceps', 'non-elbow']
MUSCLE_LABELS = {
    'biceps': 'Biceps',
    'brachioradialis': 'Brachioradialis',
    'triceps': 'Triceps',
    'non-elbow': 'Non-elbow'
}
# Color-blind friendly palette (Okabe-Ito)
MUSCLE_COLORS = {
    'biceps': '#D55E00',          # orange
    'brachioradialis': '#0072B2', # blue
    'triceps': '#CC79A7',         # pinkish purple
    'non-elbow': '#999999'        # grey
}

# ------------------ Original Circuit (without GMA lower bound) ------------------
class ExtensorCircuitWithDepression:
    def __init__(self, R=12.0, base_decay=0.1, min_decay=0.01, max_gma=20):
        self.R = R
        self.base_decay = base_decay
        self.min_decay = min_decay
        self.max_gma = max_gma
        self.target_size = 100

        # Neuron pools: brachioradialis force = -6 (moderate flexor)
        self.neuron_props = {
            'flex_strong':  {'count': 100, 'force': -9},
            'flex_mod':     {'count': 100, 'force': -6},
            'non_elbow':    {'count': 5000, 'force': 0},
            'extensor':     {'count': 100, 'force': 9}
        }
        self._init_pools()
        self.history = {'force': [], 'gma': [], 'extensor_ratio': [],
                        'reinforced_composition': []}

    def _init_pools(self):
        self.all_neurons = []
        for p in self.neuron_props.values():
            self.all_neurons.extend([p['force']] * p['count'])
        self.reinforced = random.sample(self.all_neurons, self.target_size)
        self.non_reinforced = self._subtract(self.all_neurons, self.reinforced)

    def _subtract(self, a, b):
        ca, cb = Counter(a), Counter(b)
        for k, v in cb.items():
            if ca[k] < v:
                raise ValueError('subset error')
        return list((ca - cb).elements())

    def _safe_remove(self, lst, to_remove):
        cnt = Counter(to_remove)
        new = []
        for x in lst:
            if cnt[x] > 0:
                cnt[x] -= 1
            else:
                new.append(x)
        return new

    def _activation(self, gma):
        spec_size = 15 + int(gma * 0.01 * len(self.reinforced))
        spec_size = min(spec_size, len(self.reinforced))
        spec = random.sample(self.reinforced, spec_size) if spec_size > 0 else []
        if gma > 0:
            nonspec_size = int(gma * 0.001 * len(self.non_reinforced))
            if nonspec_size > 0 and len(self.non_reinforced) > 0:
                nonspec_size = min(nonspec_size, len(self.non_reinforced))
                nonspec = random.sample(self.non_reinforced, nonspec_size)
            else:
                nonspec = []
        else:
            nonspec = []
        return spec + nonspec

    def _reinforce(self, activated):
        self.reinforced.extend(activated)
        self.non_reinforced = self._safe_remove(self.non_reinforced, activated)

    def _depress(self, activated, gma):
        rate = self.min_decay + (gma / self.max_gma) * (self.base_decay - self.min_decay)
        active_in_reinf = [x for x in activated if x in self.reinforced]
        n_remove = int(len(active_in_reinf) * rate)
        if n_remove > 0 and active_in_reinf:
            max_removable = max(0, len(self.reinforced) - 1)
            n_remove = min(n_remove, max_removable, len(active_in_reinf))
            if n_remove > 0:
                to_remove = random.sample(active_in_reinf, n_remove)
                self.reinforced = self._safe_remove(self.reinforced, to_remove)
                self.non_reinforced.extend(to_remove)

    def _prune(self):
        if len(self.reinforced) > self.target_size:
            excess = len(self.reinforced) - self.target_size
            indices_to_remove = random.sample(range(len(self.reinforced)), excess)
            for idx in sorted(indices_to_remove, reverse=True):
                pruned = self.reinforced.pop(idx)
                self.non_reinforced.append(pruned)

    def simulate(self, max_trials=500):
        gma = 0
        consec = 0
        stab_step = None
        for step in range(max_trials):
            activated = self._activation(gma)
            F = sum(activated)

            total = len(self.reinforced)
            if total == 0:
                step_composition = {muscle: 0.0 for muscle in FORCE_TO_MUSCLE.values()}
            else:
                comp = Counter(self.reinforced)
                step_composition = {muscle: comp.get(force, 0) / total
                                     for force, muscle in FORCE_TO_MUSCLE.items()}
            self.history['reinforced_composition'].append(step_composition)

            if F > self.R:
                self._reinforce(activated)
                self._prune()
                consec += 1
                gma = max(0, gma - 2)          # can go down to 0
                if consec >= 10 and stab_step is None:
                    stab_step = step
            else:
                self._depress(activated, gma)
                consec = 0
                gma = min(gma + 1, self.max_gma)

            self.history['force'].append(F)
            self.history['gma'].append(gma)
            ext_count = sum(1 for x in self.reinforced if x == 9)
            ext_ratio = ext_count / len(self.reinforced) if self.reinforced else 0
            self.history['extensor_ratio'].append(ext_ratio)

        return stab_step, self.history


# ------------------ Modified Circuit with GMA lower bound after step 400 ------------------
class ExtensorCircuitWithGMALowerBound(ExtensorCircuitWithDepression):
    def __init__(self, R=12.0, base_decay=0.1, min_decay=0.01, max_gma=20,
                 gma_lower_bound=10, constraint_start=400):
        super().__init__(R, base_decay, min_decay, max_gma)
        self.gma_lower_bound = gma_lower_bound
        self.constraint_start = constraint_start

    def simulate(self, max_trials=500):
        gma = 0
        consec = 0
        stab_step = None
        for step in range(max_trials):
            activated = self._activation(gma)
            F = sum(activated)

            total = len(self.reinforced)
            if total == 0:
                step_composition = {muscle: 0.0 for muscle in FORCE_TO_MUSCLE.values()}
            else:
                comp = Counter(self.reinforced)
                step_composition = {muscle: comp.get(force, 0) / total
                                     for force, muscle in FORCE_TO_MUSCLE.items()}
            self.history['reinforced_composition'].append(step_composition)

            if F > self.R:
                self._reinforce(activated)
                self._prune()
                consec += 1
                # After constraint_start, prevent GMA from dropping below gma_lower_bound
                if step >= self.constraint_start:
                    gma = max(gma - 2, self.gma_lower_bound)
                else:
                    gma = max(0, gma - 2)
                if consec >= 10 and stab_step is None:
                    stab_step = step
            else:
                self._depress(activated, gma)
                consec = 0
                gma = min(gma + 1, self.max_gma)

            self.history['force'].append(F)
            self.history['gma'].append(gma)
            ext_count = sum(1 for x in self.reinforced if x == 9)
            ext_ratio = ext_count / len(self.reinforced) if self.reinforced else 0
            self.history['extensor_ratio'].append(ext_ratio)

        return stab_step, self.history


# ------------------ Batch simulation (original) ------------------
def run_experiment_original(R, n_runs=20, max_trials=500):
    all_compositions = []
    all_force_histories = []
    all_gma_histories = []
    stable_count = 0
    stab_trials = []

    for run in range(n_runs):
        model = ExtensorCircuitWithDepression(R=R, base_decay=0.1, min_decay=0.01)
        sf, hist = model.simulate(max_trials)
        all_compositions.append(hist['reinforced_composition'])
        all_force_histories.append(hist['force'])
        all_gma_histories.append(hist['gma'])
        if sf is not None:
            stable_count += 1
            stab_trials.append(sf)

    if stab_trials:
        avg_step = np.mean(stab_trials)
        print(f"Original R={R}: {stable_count}/{n_runs} stabilized, avg step {avg_step:.1f}")
    else:
        print(f"Original R={R}: {stable_count}/{n_runs} stabilized")

    n_muscles = len(MUSCLE_ORDER)
    data = np.zeros((n_runs, max_trials, n_muscles))
    for r, comp_list in enumerate(all_compositions):
        for t, comp_dict in enumerate(comp_list):
            for m_idx, muscle in enumerate(MUSCLE_ORDER):
                data[r, t, m_idx] = comp_dict.get(muscle, 0.0)

    mean = np.mean(data, axis=0)
    sem = stats.sem(data, axis=0)
    ci = sem * stats.t.ppf(0.975, n_runs - 1)

    trials = list(range(max_trials))
    stats_dict = {}
    for m_idx, muscle in enumerate(MUSCLE_ORDER):
        stats_dict[muscle] = {
            'mean': mean[:, m_idx],
            'ci': ci[:, m_idx]
        }

    return trials, stats_dict, all_compositions, all_force_histories, all_gma_histories


# ------------------ Batch simulation (modified with GMA lower bound) ------------------
def run_experiment_modified(R, n_runs=20, max_trials=500,
                            gma_lower_bound=10, constraint_start=400):
    all_compositions = []
    all_force_histories = []
    all_gma_histories = []
    stable_count = 0
    stab_trials = []

    for run in range(n_runs):
        model = ExtensorCircuitWithGMALowerBound(R=R, base_decay=0.1, min_decay=0.01,
                                                  gma_lower_bound=gma_lower_bound,
                                                  constraint_start=constraint_start)
        sf, hist = model.simulate(max_trials)
        all_compositions.append(hist['reinforced_composition'])
        all_force_histories.append(hist['force'])
        all_gma_histories.append(hist['gma'])
        if sf is not None:
            stable_count += 1
            stab_trials.append(sf)

    if stab_trials:
        avg_step = np.mean(stab_trials)
        print(f"Modified R={R} (lower bound={gma_lower_bound} after {constraint_start}): {stable_count}/{n_runs} stabilized, avg step {avg_step:.1f}")
    else:
        print(f"Modified R={R} (lower bound={gma_lower_bound} after {constraint_start}): {stable_count}/{n_runs} stabilized")

    n_muscles = len(MUSCLE_ORDER)
    data = np.zeros((n_runs, max_trials, n_muscles))
    for r, comp_list in enumerate(all_compositions):
        for t, comp_dict in enumerate(comp_list):
            for m_idx, muscle in enumerate(MUSCLE_ORDER):
                data[r, t, m_idx] = comp_dict.get(muscle, 0.0)

    mean = np.mean(data, axis=0)
    sem = stats.sem(data, axis=0)
    ci = sem * stats.t.ppf(0.975, n_runs - 1)

    trials = list(range(max_trials))
    stats_dict = {}
    for m_idx, muscle in enumerate(MUSCLE_ORDER):
        stats_dict[muscle] = {
            'mean': mean[:, m_idx],
            'ci': ci[:, m_idx]
        }

    return trials, stats_dict, all_compositions, all_force_histories, all_gma_histories


# ------------------ Plotting function for comparison ------------------
def plot_comparison(stats_orig, stats_mod, output_base='Extensor_R2_Comparison_GMA_LowerBound'):
    """Compare original R=2 vs modified R=2 (GMA lower bound after 400)"""
    def smooth_trace(data, window=9):
        if len(data) < window:
            return data
        kernel = np.ones(window) / window
        smoothed = np.convolve(data, kernel, mode='same')
        pad = window // 2
        smoothed[:pad] = data[:pad]
        smoothed[-pad:] = data[-pad:]
        return smoothed

    plt.rcParams.update({
        'font.family': 'sans-serif',
        'font.sans-serif': ['Arial', 'Helvetica'],
        'font.size': 7,
        'axes.labelsize': 8,
        'axes.titlesize': 9,
        'axes.spines.top': False,
        'axes.spines.right': False,
        'axes.linewidth': 0.8,
        'xtick.direction': 'in',
        'ytick.direction': 'in',
        'legend.fontsize': 6,
        'figure.dpi': 300,
        'savefig.dpi': 300,
        'savefig.bbox': 'tight'
    })

    fig, axes = plt.subplots(2, 2, figsize=(7.2, 5.5))
    # Row 0: Original R=2, Row 1: Modified R=2
    # Each row: left = extensor_ratio, right = GMA

    # Color definitions matching original Fig2A.py
    color_gma = '#ff7f0e'   # Orange, consistent with original GMA color
    color_lower_bound = 'red'

    for row, (stats_data, label) in enumerate(zip([stats_orig, stats_mod],
                                                   ['Original R=2', 'Modified R=2 (lower bound 10 after 400)'])):
        trials, stats_dict, comps, forces, gmas = stats_data

        # ---- Left: extensor_ratio (triceps proportion) ----
        ax_ratio = axes[row, 0]
        ext_mean = stats_dict['triceps']['mean']
        ext_ci = stats_dict['triceps']['ci']
        ext_mean_smooth = smooth_trace(ext_mean, window=9)
        ax_ratio.plot(trials, ext_mean_smooth, color=MUSCLE_COLORS['triceps'], lw=1.5, label='Extensor (triceps) proportion')
        ax_ratio.fill_between(trials, ext_mean - ext_ci, ext_mean + ext_ci,
                              color=MUSCLE_COLORS['triceps'], alpha=0.2)
        ax_ratio.axvline(x=400, color='gray', linestyle='--', lw=1, label='Constraint onset')
        ax_ratio.set_xlabel('Time (trials)')
        ax_ratio.set_ylabel('Extensor proportion in reinforced pool')
        ax_ratio.set_title(label, fontweight='bold')
        ax_ratio.set_ylim(0, 1)
        ax_ratio.spines['top'].set_visible(False)
        ax_ratio.spines['right'].set_visible(False)

        # ---- Right: GMA (mean ± CI) ----
        ax_gma = axes[row, 1]
        gma_array = np.array([h[:len(trials)] for h in gmas])
        g_mean = np.mean(gma_array, axis=0)
        g_sem = stats.sem(gma_array, axis=0)
        g_ci = g_sem * stats.t.ppf(0.975, gma_array.shape[0] - 1)

        g_mean_smooth = smooth_trace(g_mean, window=9)
        ax_gma.plot(trials, g_mean_smooth, color=color_gma, lw=1.5, label='GMA')
        ax_gma.fill_between(trials, g_mean - g_ci, g_mean + g_ci,
                            color=color_gma, alpha=0.2)
        ax_gma.axvline(x=400, color='gray', linestyle='--', lw=1)
        ax_gma.axhline(y=10, color=color_lower_bound, linestyle=':', lw=1, label='GMA lower bound')
        ax_gma.set_xlabel('Time (trials)')
        ax_gma.set_ylabel('Global Modulatory Activation (GMA)')
        ax_gma.set_title('GMA dynamics', fontweight='bold')
        ax_gma.set_ylim(0, 21)
        ax_gma.spines['top'].set_visible(False)
        ax_gma.spines['right'].set_visible(False)

        # legends
        ax_ratio.legend(loc='upper right', frameon=False, fontsize=6)
        ax_gma.legend(loc='upper right', frameon=False, fontsize=6)

    plt.tight_layout()
    for ext in ['pdf', 'svg']:
        plt.savefig(f"{output_base}.{ext}", dpi=300, bbox_inches='tight')
    plt.close()
    print(f"Saved {output_base}.pdf and .svg")


# ------------------ Main ------------------
if __name__ == "__main__":
    random.seed(32)
    np.random.seed(32)

    # Run original R=2
    print("Running original R=2 ...")
    stats_orig_R2 = run_experiment_original(R=2, n_runs=20, max_trials=500)

    # Run modified R=2 with GMA lower bound after 400
    print("\nRunning modified R=2 (lower bound=10 after step 400) ...")
    stats_mod_R2 = run_experiment_modified(R=2, n_runs=20, max_trials=500,
                                           gma_lower_bound=10, constraint_start=400)

    # Plot comparison
    plot_comparison(stats_orig_R2, stats_mod_R2,
                    output_base='D Extensor_R2_Comparison_GMA_LowerBound')
```
